# Multiparameter optimization extends the lifetime of cell-free protein synthesis in a high-throughput format

**DOI:** 10.64898/2026.07.15.738603

**Authors:** Eray U. Bozkurt, Baptiste Zanchet, Pablo I. Nikel, Daniel C. Volke

## Abstract

Cell-free protein synthesis (CFPS) is a powerful platform for synthetic biology, yet the factors governing reaction longevity remain poorly understood despite their importance for high-throughput applications. Here, the three principal determinants of CFPS performance—DNA template design, reaction composition, and lysate genotype—were systematically optimized to extend reaction lifetime in a 384-well plate format. Different energy regeneration systems were evaluated through real-time pH monitoring and metabolomic analyses to identify the metabolic constraints limiting prolonged protein synthesis. Lysates prepared from engineered *Escherichia coli* BL21(DE3) strains were further examined to assess the contributions of DNA, RNA, and amino acid stabilization. Systematic optimization of amino acid, nucleoside triphosphate, polyethylene glycol, and lysate concentrations identified DNA template stability and amino acid preservation as the primary factors sustaining CFPS activity. Combining these improvements yielded reactions that remained productive for >14 h and produced 567 ± 64 μg mL^−1^ active deGFP. These findings establish practical strategies for extending CFPS lifetime and improving high-throughput cell-free platforms.

## Introduction

Cell-free protein synthesis (CFPS) emerged in the early 1960s from efforts to decipher the genetic code.^1, 2^ These pioneering studies demonstrated that complex biological processes can be reconstituted outside living cells.^3^ The rise of synthetic biology has since expanded the scope of cell-free systems well beyond mechanistic biochemistry and molecular biology. CFPS is now used for genetic part screening,^4–6^ incorporation of non-canonical amino acids,^7–9^ pathway prototyping,^10–14^ biosensing and diagnostics,^15–18^ and production of toxic (or otherwise difficult to synthesize *in vivo*) proteins,^19^ among many other applications. Entire viral particles have even been assembled *in vitro* in cell-free systems.^20^ This broad adoption increased the need to understand the biochemical and physical constraints that determine CFPS performance.^21, 22^

CFPS relies on three core components:^3^ the cell lysate, which provides the transcription and translation machinery; the reaction buffer, which supplies metabolites, ions, and energy; and the DNA template, which encodes the protein of interest. Each component has been refined over decades of CFPS research.^23^ Early systems relied on high-energy phosphate donors (e.g., phosphoenolpyruvate), but these metabolites caused rapid accumulation of inorganic phosphate, thereby limiting or even inhibiting protein synthesis. Later studies showed that glycolysis remains active in crude lysates, enabling glucose,^24^ maltodextrin, and maltose^25^ to regenerate ATP with higher efficiency and lower cost. The energy source also shapes the pH trajectory of the CFPS reaction. Glucose can drive rapid acidification,^26^ which must be counteracted by strong buffering systems.^27^ By contrast, maltodextrin and related substrates support more stable metabolic fluxes that can sustain CFPS.

Macromolecular crowding is another critical determinant of the overall CFPS performance. The *Escherichia coli* cytoplasm, for instance, is densely packed with proteins, nucleic acids, and metabolites, and this crowded environment strongly influences enzyme activity, molecular diffusion, and reaction kinetics.^28, 29^ Because lysates are typically diluted ca. 20-fold during reaction assembly, they do not fully reproduce the intracellular bacterial environment.^24, 30, 31^ Molecular crowding agents, e.g., polyethylene glycol (PEG), Ficoll, or dextran, are therefore added to restore excluded volume effects and mimic physiological conditions.^32, 33^

Lysate composition also has a major impact on reaction performance. *E. coli* is the dominant lysate source because of its expression capacity and extensively characterized proteome. *E. coli* BL21 (DE3) is widely used because it encodes the T7 RNA polymerase (T7 RNAP) and lacks several proteases that would otherwise compromise protein stability.^3^ To enhance performance further, genetic engineering of lysate donor strains has been implemented to stabilize DNA,^34^ RNA,^35^ amino acids,^36^ and recombinant proteins,^37^ and to promote incorporation of non-canonical amino acids.^7^ These efforts have primarily focused on increasing protein yield, whereas their effects on CFPS dynamics and reaction longevity remain much less explored.^38, 39^

Here, we systematically evaluated the three central components of CFPS in 384-well plates compatible with high-throughput formats. DNA template design, buffer composition, and lysate donor genotype were analyzed to identify conditions that support extended protein synthesis (**Figure 1A**). Different energy sources were compared, and their effects on reaction pH and metabolite dynamics were thoroughly characterized with dedicated sensors and omic approaches. Furthermore, lysates from engineered *E. coli* BL21 (DE3) were evaluated to determine their effects on reaction duration. Amino acids, nucleoside triphosphates, and PEG were titrated to define their contributions to reaction longevity. Unlike previous studies that primarily focused on endpoint protein yields and single-parameter optimization, this work integrates lysate engineering, metabolic characterization, and reaction optimization to identify factors governing CFPS longevity in a high-throughput microplate format. Collectively, these analyses establish a framework for developing long-duration CFPS reactions compatible with high-throughput screening platforms.

**Figure 1.**
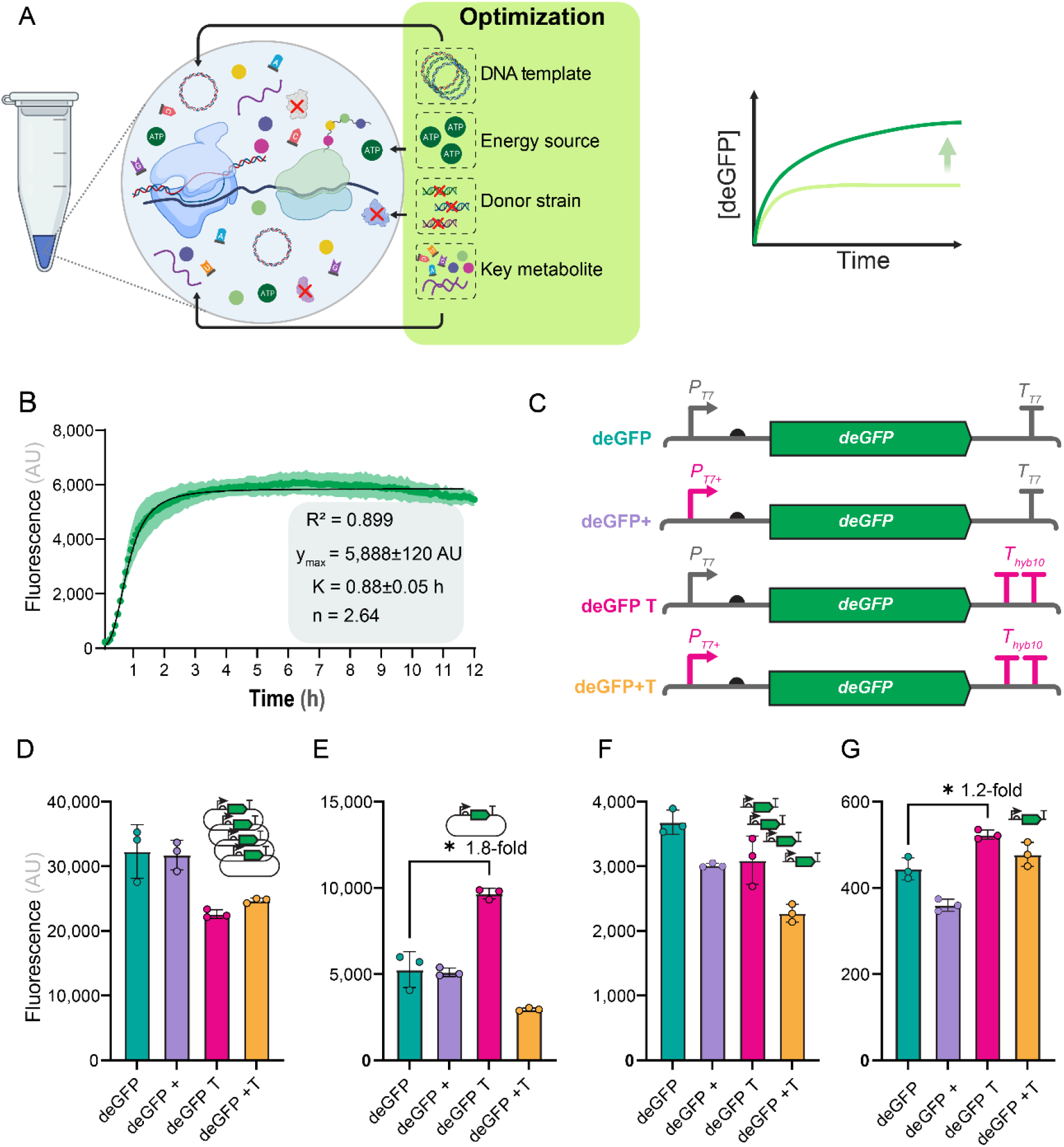
Benchmarking initial parameters for high-throughput cell-free protein synthesis (CFPS) reactions. **(A)** Overview of the core components of the CFPS system and the parameters evaluated. **(B)** GFP production over time in CFPS reactions used to assess reproducibility. Parameters obtained from fitting the data to **Equation 1** are shown in the inset. Plasmid pT7·GFPmut3-TT7 was used as the expression template. **(C)** Architecture of plasmid and linear DNA templates containing engineered promoter and terminator variants. Plasmid templates were evaluated in replicated CFPS reactions at high **(D)** (10 nM) and low **(E)** template concentrations (1 nM). Linear expression templates (LETs) were tested under the same conditions at high **(F)** (25 nM) and low **(G)** concentrations (2.5 nM). Data represent mean values ± standard deviation of three experiments performed from independently prepared lysate batches. Technical replicates were averaged before statistical analysis. Details of the statistical analyses and curve-fitting parameters are provided in **File S1**.

## Results and Discussion

### Establishing Baseline Conditions and Template Engineering for CFPS

Developing a multiparameter screening strategy for CFPS requires a reaction setup that allows systematic variation of individual components without introducing unnecessary experimental variability. For this reason, NTPs and energy sources were supplied separately to the reaction mixture, as both components were later tested across different concentrations. Because lysates from engineered genetic backgrounds were also evaluated, high reproducibility was essential. CFPS reactions can vary substantially because of their biochemical complexity and the many parameters that influence reaction output.^40^ Initial experiments therefore focused on establishing baseline performance by assessing batch-to-batch reproducibility, defining the lysate-to-buffer ratio, and identifying a suitable DNA template for protein expression.

Four lysate batches were prepared on separate days to evaluate reproducibility. GFPmut3 production from plasmid pT7·GFPmut3-TT7 was used as the readout of lysate performance. The four lysates produced highly consistent endpoint fluorescence values, with less than 13.2% inter-day variation (**Figure 1B**). The CFPS reaction displayed a characteristic sigmoidal protein accumulation profile, consisting of an initial lag phase, a rapid production phase, and a terminal plateau. The fluorescence trajectory was therefore analyzed using a Hill-type sigmoidal model to estimate reaction yield (*y*_max)_, reaction half-time (*K*), and curve steepness (*n*). Interpreting the CFPS output fluorescence with **Equation 1** provided an excellent fit to the data (*R*^2^ = 0.899), with *K* = 0.88 ± 0.05 h and *y*_max =_ 5,888 ± 120 arbitrary units (AU).

After validating the system and establishing a benchmark for further optimization, we examined the effect of DNA template design on protein synthesis. Three plasmids encoding different GFP variants (i.e., msfGFP^41^, GFPmut3,^42^ and deGFP^43^) were compared to this end. Their architectures and general characteristics are shown in **Figure S2**. msfGFP is a monomeric superfolder GFP optimized for stable folding and contains additional mutations that increases the fluorescence output. GFPmut3 was optimized for fluorescence-assisted cell sorting and carries two mutations close to the chromophore that increase brightness. deGFP is derived from eGFP and was improved for CFPS by deleting the first 6 amino acids and removing RBS-like structures from the open reading frame. The deGFP construct containing a dedicated T7 terminator produced the highest protein levels in our system (**Figure S2**), consistent with previous observations that *deGFP* is translated more efficiently in CFPS than other GFP variants.^43^

Promoter and terminator sequences were then modified to test whether transcriptional or translational features limited protein synthesis by generating three additional variants (**Figure 1C**). A construct carrying a transcription-enhancing sequence at the +4 site of the promoter region, 5’-AAATA-3’ instead of 5’-AGACC-3’, previously reported to increase T7 promoter activity,^44^ was designated deGFP+. In the second construct, the T7 terminator was replaced with an engineered double terminator, TT7*_hyb10_*,^45^ and the construct was designated deGFP T. This strong terminator prevents readthrough by T7 RNAP and can also stabilize the mRNA. The third construct, combining both elements, was designated deGFP+T. Our experiments indicated that neither modification increased deGFP production in CFPS (**Figure 1D**). Thus, transcriptional output and template architecture did not appear to be the primary limiting factors under the tested conditions. In these reactions, the circular DNA template was supplied at a relatively high concentration (10 nM). We reasoned that limiting template availability could impose a greater demand on transcriptional and translational efficiency; hence, we repeated the experiments with 1 nM circular DNA as the template (**Figure 1E**). Under this condition, the deGFP T construct produced significantly more protein (1.8-fold), probably because increased mRNA stability allowed more translation.^46^

To evaluate this behavior further, linear expression templates (LETs) were generated for each construct. LETs are more susceptible to degradation and therefore impose stronger constraints on template stability. LETs were amplified with ca. 300 bp of flanking sequence on each terminus and supplied to CFPS reactions at high DNA concentration (25 nM, **Figure 1F**) or limiting concentration (2.5 nM, **Figure 1G**). Consistent with the observations obtained using plasmid-based templates, deGFP T performed significantly better at low DNA concentration (1.2-fold), whereas no difference was observed at high DNA concentration. The transcription-enhancing promoter in the deGFP+ construct, however, did not improve deGFP yield under any tested condition.

Taken together, these findings indicate that transcript stability becomes a defining factor under low DNA availability. This parameter is relevant for high-throughput screening formats that rely on low template loads, where mRNA stability can have a greater influence than transcriptional output. By contrast, transcription does not appear to be limiting under the tested conditions, reflecting the high activity of T7 RNAP and the absence of competing recruitment processes. Because subsequent experiments used higher DNA template amounts, the standard template without promoter or terminator modification was selected for the rest of the study.

### Selecting an Optimal Energy Source for Prolonged CFPS Performance

CFPS reactions depend on external metabolites for ATP regeneration. Protein synthesis is energy intensive, and ATP is consumed at multiple steps.^47^ Several metabolic routes have been used to support ATP regeneration in a cost-effective and metabolically efficient manner. We therefore evaluated five low-cost energy sources, i.e., ribose, glycogen, glucose, maltodextrin, and maltose, across different concentrations to assess their effects on protein yield and reaction duration.

Glucose is commonly used in CFPS, and this sugar is metabolized rapidly by crude lysates. Rapid sugar utilization is likely linked to the high abundance of glycolytic enzymes in the extract, as the lysate donor strain was grown with glucose as the main carbon source. However, fast glucose metabolism can also promote accumulation of inhibitory metabolites. Energy sources that are metabolized more slowly may alleviate this limitation. Glycogen and maltodextrin are glucose polymers that enter glycolysis after depolymerization (**Figure 2A**). This additional depolymerization step slows the overall rate of substrate utilization. In addition, phosphorylation during polymer breakdown uses inorganic phosphate rather than ATP, which improves energy efficiency and recycles inorganic phosphate generated during CFPS. Both polymers are synthesized by *E*. *coli* as carbon and energy reserves, and enzymes for their degradation are constitutively present in cells.^48^ Maltose is a glucose dimer that is converted into glucose and glucose-6-phosphate through inorganic phosphate-dependent phosphorylation. Ribose is activated by ATP-dependent phosphorylation, as is glucose, and then enters metabolism through the pentose phosphate pathway. Because flux through the pentose phosphate pathway is generally lower than glycolytic flux in *E*. *coli*, slower ribose utilization was anticipated.

**Figure 2.**
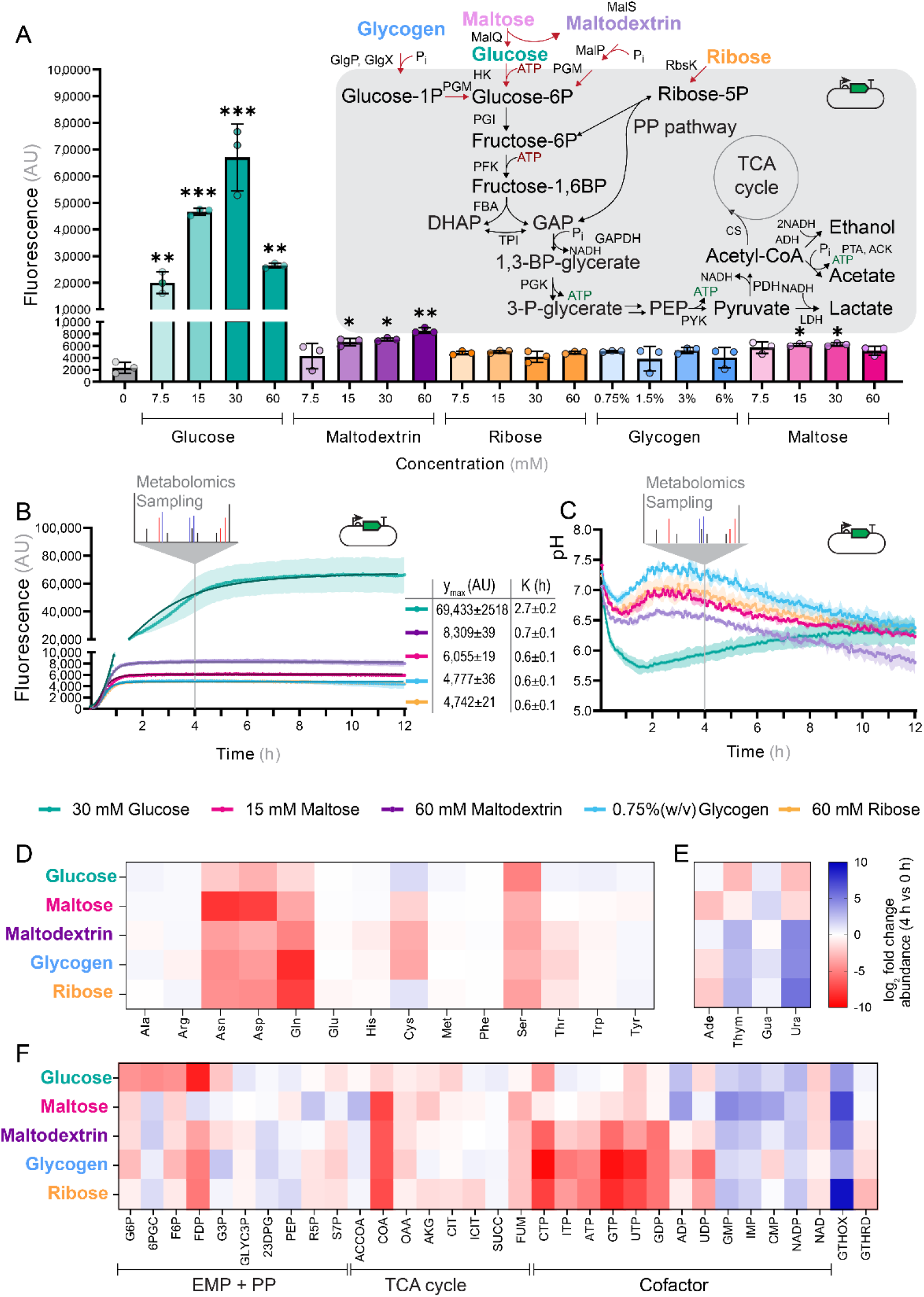
Influence of energy sources on CFPS performance. **(A)** Five energy sources were tested at different concentrations to fuel CFPS reactions for deGFP production. **(B)** Time course of GFP synthesis using the optimal concentration of each energy source. The metabolomics sampling point after 4 h is indicated. **(C)** Real-time pH measurements obtained with purified PHP in reactions containing different energy sources. Changes in amino acid pools **(D)**, nucleosides **(E)**, and other metabolites **(F)** after 4 h are indicated relative to initial levels. Metabolites and enzyme abbreviations follow the BiGG Models database. PP, pentose phosphate; TCA cycle, tricarboxylic acid cycle. Data in panels **(A–C)** represent mean values ± standard deviation from three experiments performed with independently prepared lysate batches. Technical replicates were averaged before statistical analysis. Data in panels **(D–F)** represent mean values from three pooled replicates prepared from independently generated lysate batches. Details of the statistical analyses and curve-fitting parameters are provided in **File S1**.

All tested energy sources increased mean protein production relative to the control (**Figure 2A**). Glucose, maltodextrin, and ribose significantly increased protein production at several concentrations, showing that these substrates can fuel CFPS. Reactions supplemented with 30 mM glucose produced the highest amount of protein, with a 28-fold increase relative to the control. The optimal concentrations for the other substrates were 15 mM ribose, yielding a 2.2-fold increase; 3% (w/v) glycogen, yielding a 2.2-fold increase; 60 mM maltodextrin, yielding a 3.6-fold increase; and 30 mM maltose, yielding a 2.7-fold increase. Reaction times were then quantified using the best concentration for each substrate (**Figure 2B**). CFPS reactions fueled by all compounds except glucose reached completion at similar times, with *K* = 0.6-0.7 h. By contrast, CFPS reactions fueled by glucose displayed biphasic production kinetics and a substantially extended production phase, with *K* = 2.7 ± 0.2 h.

Reaction pH is a major determinant of CFPS duration and efficiency, and pH 6.8 has been reported as optimal for driving protein expression in CFPS systems.^27^ To monitor pH dynamics during CFPS, we used the pH-sensitive GFP variant PHP for real-time pH tracking.^49^ This variant had previously been used as a pH sensor in the cytoplasm of different bacterial species, including *E*. *coli*. To minimize changes to the CFPS setup and avoid interference between newly synthesized GFP and the pH sensor, we generated a non-fluorescent GFP variant, dGFP, by introducing several amino acid substitutions that suppress fluorescence. CFPS reactions were then assembled with a DNA template encoding dGFP, and purified PHP protein was added to each reaction. Only reactions containing glucose showed a rapid and pronounced pH decrease (**Figure 2C**). This result agrees with previous findings showing that glucose is utilized rapidly and that this metabolism is accompanied by acid formation, leading to a sharp decrease in pH.^26^ This behavior may reflect the abundance of glycolytic enzymes in the lysate, as the donor *E*. *coli* strain was cultivated with glucose as the primary carbon and energy source. For the other energy sources, only modest acidification was observed during the first hour, followed by partial pH recovery. The acidification period coincided with the timing and intensity of protein synthesis. Fast glucose utilization likely provided sufficient energy to saturate transcription and translation, thereby maximizing protein synthesis, but also caused accumulation of organic acids as by-products. By contrast, the other energy sources appeared to be metabolized too slowly to fully sustain gene expression. The marked reduction in protein synthesis after ca. 1 h across all energy sources suggests that additional factors limit CFPS activity. For the conditions where glucose acted as the main carbon substrate, acidification may further slow protein synthesis.

We next assessed how each energy source affected the cell-free metabolome. Metabolites were quantified at the start of the reaction and after 4 h, when protein synthesis had ceased. Amino acids showed similar depletion patterns across all conditions (**Figure 2D**). Asparagine, aspartate, glutamine, and serine decreased sharply with all energy sources. Because the lysate donor strain was grown in rich medium, where amino acids are supplied in excess, amino acid degradation pathways were likely induced and the corresponding enzymes retained in the CFPS reaction.^50^ Previous work showed that serine, aspartate, and glutamate are consumed at rates that typically exceed their incorporation into protein.^51^ Glutamine can also serve as an amine donor in transaminase reactions and may therefore be consumed independently of protein synthesis. Notably, amino acid depletion was generally lower when glucose was used as the energy source, possibly indicating that rapid glucose metabolism contributed to replenishment of amino acid pools. This general effect could help explain the longer productive duration of glucose-fueled reactions.

Nucleobase levels showed a substrate-dependent pattern. Thymine and uracil were depleted with glucose and, to a lesser extent, with maltose, whereas their concentrations substantially increased in the presence of the other energy sources (**Figure 2E**). Nucleobases are building blocks for nucleotides, which are required for supporting transcription. This result is consistent with previous reports showing that nucleotides can be efficiently synthesized in CFPS.^52^ The increase in nucleobases observed with the other substrates likely reflects nucleotide degradation rather than sustained nucleotide biosynthesis.

Glucose also stabilized nucleotide triphosphate concentrations (**Figure 2F**). CFPS reactions with maltose retained higher nucleotide triphosphate levels as well. This observation indicates that energy supply continued after 4 h in reactions fueled by glucose or maltose, whereas catabolism of the other energy sources had already slowed or stopped. Glucose also caused clear differences in glycolytic and tricarboxylic acid (TCA) cycle metabolites. Maltodextrin, glycogen, and maltose did not energize CFPS as effectively as glucose, even though these substrates are ultimately converted into glucose-6-phosphate and enter the same metabolic routes. The utilization of these substrates depends on initial phosphorylation using inorganic phosphate. Although inorganic phosphate is theoretically recycled in the reaction, an imbalance in phosphate availability may interrupt polymer breakdown and limit ATP regeneration.

Although glucose supported the highest yields and sustained energy regeneration, rapid sugar metabolism caused strong acidification of the cell-free milieu. Using glucose would therefore require stronger buffering formulations than those applied here, which may come at the price of inhibiting or slowing protein synthesis. Slower-metabolizing substrates produced comparable reaction kinetics and more stable pH trajectories. Among these substrates, maltodextrin showed the strongest performance and was selected as the energy source for subsequent assays. Because protein synthesis appeared to be limited by additional factors, e.g., amino acid depletion, we next modified lysate metabolism to support extended reaction times.

### Genome Engineering of the Lysate donor to Support Efficient CFPS

*E. coli* BL21 (DE3) is widely used for lysate preparation due to its high expression capacity through the T7 RNAP system and the absence of several proteases.^3^ Genetic modification of this strain can further improve CFPS for defined applications, since many native enzymes are predicted to exert a detrimental effect on reaction performance.^53^ We therefore engineered *E. coli* BL21 (DE3) to stabilize DNA (Δ*endA*), RNA (Δ*rnB* Δ*rnE* Δ*mazF* Δ*csdA*), redox metabolism (Δ*gorA*) **(Figure 3A)**, and amino acids (ΔAA; i.e., Δ*gshA* Δ*tnaA* Δ*speA* Δ*sdaA* Δ*sdaB*) **(Figure 3B)**.^7, 54, 55^ EndA is a periplasmic endonuclease, and its elimination should stabilize plasmid DNA; for this reason, *endA* inactivation is a common modification in cloning *E*. *coli* strains.^56^ CsdA is a cold-shock-responsive RNA helicase that forms a degradome with ribonuclease RNasE upon cold exposure.^57, 58^ MazF is an endoribonuclease and part of a toxin-antitoxin system, while RNase II (encoded by *rnB*) is an exoribonuclease.^59^ DNA and RNA stability are major determinants of CFPS efficiency, and these modifications have been previously shown to positively influence protein yields.^60^ Glutathione reductase, encoded by *gorA*, consumes reducing equivalents to maintain redox homeostasis, a function that is not required in CFPS.^53^ Glutamate-cysteine ligase (GshA), tryptophanase (TnaA), arginine decarboxylase (SpeA), serine deaminase (SdaB), and serine ammonia-lyase (SdaA) modify amino acids and can thereby deplete amino acid pools during CFPS (**Table S4**).^53, 54^ In all cases, a cytidine base-editor^55^ was used to insert premature *STOP* codons into the targeted genes,^61^ generating truncated, non-functional proteins (**Figure 3C**). This strategy enabled rapid construction of lysate donor strains carrying one or several gene knockouts. For example, all five mutations designed to stabilize amino acid pools could be introduced simultaneously by using this methodology.

**Figure 3.**
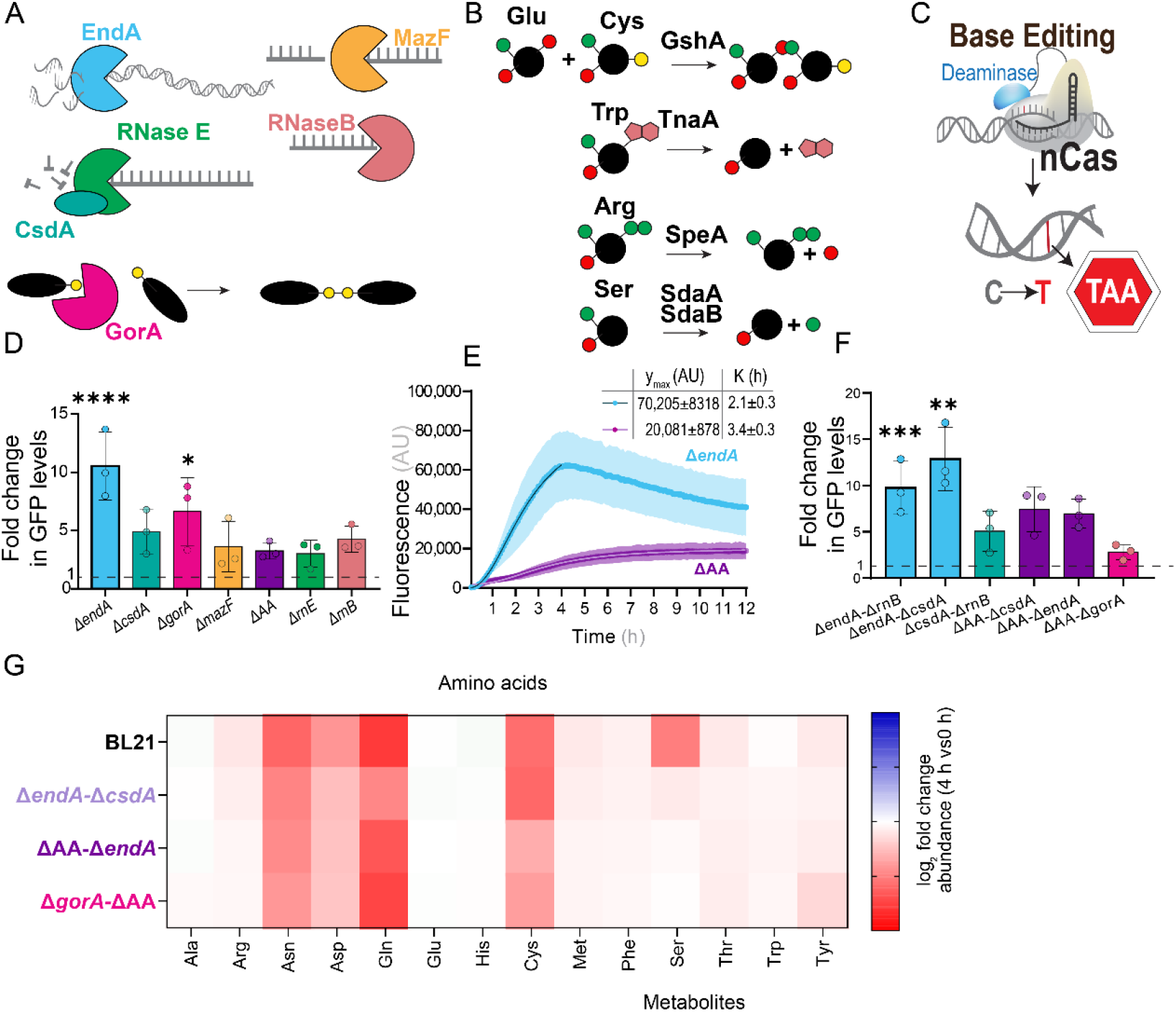
Performance of CFPS with lysates prepared from different *E. coli* mutants. **(A)** Enzymes removed from the lysate by genetic modification of the donor strain. EndA is a periplasmic endonuclease that degrades DNA. The endo-and exoribonucleases MazF, RNase E (Rne), and RNase B degrade RNA. CsdA is an RNA helicase associated with RNase E that promotes RNA degradation during cold exposure. GorA maintains glutathione in its reduced state. **(B)** Enzymatic reactions eliminated from the lysate donor strain to stabilize amino acids. Amino and carboxyl groups are shown in green and red, respectively. **(C)** Mechanism of the cytidine base-editor used to inactivate target genes. Cytidine is converted to thymine within the protospacer, enabling introduction of premature *STOP* codons into the open reading frame. **(D)** Final GFP production obtained with lysates from engineered strains compared with the parental BL21 (DE3) strain. **(E)** Time course of GFP production using lysates from the Δ*endA* and ΔAA strains. **(F)** GFP production in strains carrying combinations of deletions selected for potential synergistic effects. **(G)** Amino acid and nucleotide levels after 4 h relative to initial concentrations in GFP-producing reactions. Data in panels **(D–F)** represent mean values ± standard deviation from three experiments performed with independently prepared lysate batches. Technical replicates were averaged before statistical analysis. Data in panel **(G)** represents values from three pooled replicates prepared from independently generated lysate batches. Details of the statistical analyses and curve-fitting parameters are provided in **File S1**.

To evaluate the performance of engineered lysate donor strains, three independent lysates were prepared from each strain. CFPS reactions were incubated for 12 h, and GFP synthesis was monitored over time (**Figure 3D**). All engineered strains outperformed the parental *E*. *coli* BL21 (DE3) strain in terms of protein production: Δ*csdA*, 5.8 ± 1.9-fold; Δ*gorA*, 6.6 ± 2.9-fold; Δ*mazF*, 3.6 ± 2.2-fold; ΔAA, 3.3 ± 0.7-fold; Δ*rnE*, 3.0 ± 1.1-fold; and Δ*rnB*, 4.3 ± 1.1-fold. Disruption of *endA* increased GFP yield by 10.5 ± 2.9-fold, indicating that DNA template stability was a major determinant of CFPS performance under the tested conditions. The Δ*endA* and ΔAA lysates also displayed distinct production profiles. Lysates generated from the Δ*endA* strain showed rapid expression and reached an early maximum (*K* = 2.1 ± 0.3 h), whereas lysates from the ΔAA strain showed slower and more sustained protein accumulation (*K* = 3.4 ± 0.3 h; **Figure 3E**). These observations suggested that stabilizing amino acid pools can effectively extend reaction duration.

Additional strains were constructed to test potential synergistic effects between beneficial mutations. These combinations included Δ*endA* Δ*csdA*, Δ*endA* Δ*rnB*, Δ*csdA* Δ*rnB*, ΔAA Δ*csdA*, Δ*endA* ΔAA, and ΔAA Δ*gorA*. Combining Δ*endA* with Δ*csdA* produced the highest overall protein output (12.9 ± 3.4-fold), although this improvement was not significantly higher than that obtained with the single Δ*endA* deletion (**Figure 3F**). By contrast, the Δ*endA* ΔAA combination maintained the extended reaction profile of the amino acid stabilization strain while increasing total protein output (7.4 ± 2.4-fold; **Figure S3**).

Metabolite analysis was performed to determine whether the engineered deletions altered amino acid consumption. As expected, cysteine, serine, and arginine were less depleted in CFPS reactions prepared from strains carrying disruptions in amino acid degradation pathways (**Figure 3G**). These results substantiated that deletion of the corresponding catabolic pathways stabilizes these amino acids. Tryptophan levels decreased only slightly, even in extracts derived from the parental strain, while other amino acids showed no clear differences between lysates. Based on the combined stabilization of amino acids and DNA template, *E*. *coli* BL21 (DE3) Δ*endA* ΔAA was retained for further investigation.

### Optimization of Key Metabolites in CFPS for Extended Reactions

After selecting a lysate background that supported extended reaction duration, we systematically analyzed how key reaction components influence protein synthesis and CFPS longevity. PEG increases molecular crowding and helps approximate the cytoplasmic environment,^62^ while also increasing viscosity and reducing evaporation. Omitting PEG from the reaction resulted in poor performance, likely due to reduced encounters between DNA templates and the transcription-translation machinery (**Figure 4A**). Addition of 2% (w/v) PEG, in contrast, produced the highest protein output and the longest reaction time (*y*_max =_ 35,064 ± 555 AU and *K* = 2.4 ± 0.1 h). Higher PEG concentrations, however, reduced protein synthesis. These results are consistent with reports showing that excessive crowding can hinder enzyme and DNA dissociation, thereby slowing transcription and translation.^63^

**Figure 4.**
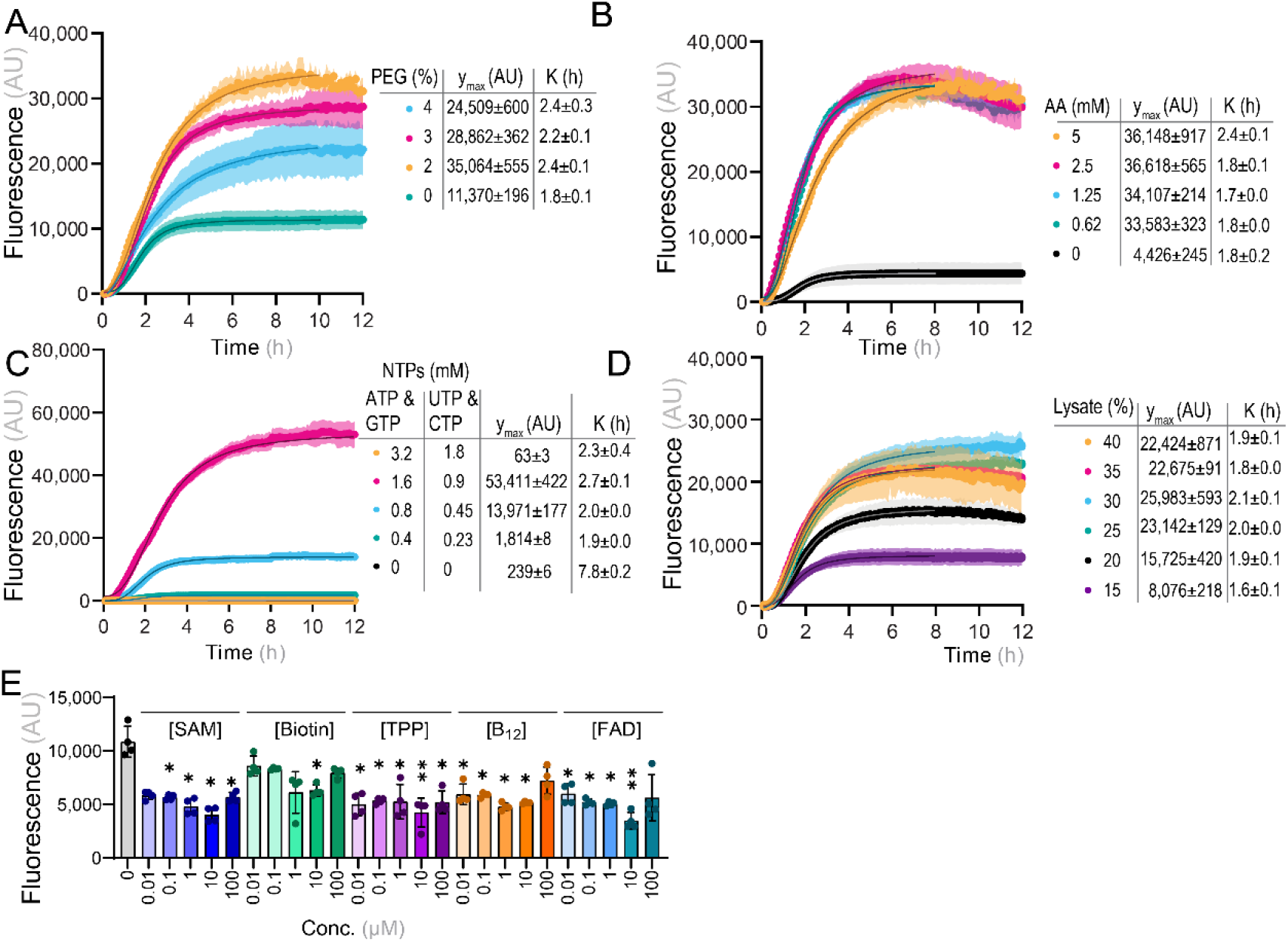
Strain-specific parameter optimization for enhanced CFPS performance. Effect of key reaction parameters on CFPS quantified by deGFP production. Polyethylene glycol (PEG) **(A)**, amino acids **(B)**, and nucleoside triphosphates (NTPs) **(C)** were titrated to determine their optimal concentrations and assess their contribution to CFPS performance. **(D)** Effect of lysate concentration on CFPS. **(E)** Effect of supplementing additional cofactors at different concentrations (concn.) on CFPS performance. Data represents mean values ± standard deviation from at least three independent experiments. Technical replicates were averaged before statistical analysis. Details of the statistical analyses and curve-fitting parameters are provided in **File S1**.

Amino acids and NTPs are the primary substrates for protein and RNA synthesis, and their concentrations strongly influence reaction efficiency. Titration of proteinogenic amino acids showed that reactions lacking supplemental amino acids produced very low protein levels (**Figure 4B**), indicating that amino acid biosynthesis in the extract is insufficient to sustain translation. Supplementation of amino acids at 0.62 mM or higher concentrations produced similar expression kinetics, although addition of 5 mM amino acids slightly extended the reaction. These results suggest that 0.62 mM amino acids is sufficient to meet translational demand under the tested conditions. Residual amino acids present in the non-dialyzed lysate likely also contributed to the observed activity levels.

NTP titration produced a distinct pattern (**Figure 4C**). Standard reactions contained 1.6 mM ATP and GTP, and 0.9 mM CTP and UTP. ATP and GTP were supplied at higher concentrations because they also act as cofactors in primary carbon metabolism, including glycolysis and the TCA cycle. Doubling the NTP concentration completely inhibited protein synthesis. This effect was not surprising, since NTPs are known to inhibit DNA primase and RNA synthesis at high concentrations and they can sequester divalent cations.^64^ In contrast, reactions lacking supplemental NTPs produced almost no protein, confirming that external NTP supply is essential.

We next examined the lysate-to-buffer ratio. The optimal lysate fraction in CFPS is often considered to be around 33% (v/v).^3^ Semi-continuous CFPS reactions have shown that protein synthesis terminates not because the transcription-translation machinery fails, but because metabolites (e.g., inorganic phosphate) accumulate and interfere with this machinery.^65^ Increasing the buffer-to-lysate ratio could therefore potentially extend reaction duration by increasing the availability of metabolites and diluting inhibitory products. To determine the optimal lysate amount, we started with 12 mg mL^−1^ lysate, corresponding to 40% of the total reaction volume, and decreased the concentration to 5.5 mg mL^−1^, corresponding to 15% of the total volume (**Figure 4D**). Reduced lysate volume was compensated with S30 buffer. Lysate concentration had only a modest effect on reaction time (*K* = 1.6 ± 0.1 h for 15% and *K* = 2.1 ± 0.1 h for 30%). Protein production rates were highest with 9 mg mL^−1^ lysate, corresponding to 30% of the reaction volume (*y*_max =_ 25,983 ± 593 AU). Lysate concentrations below 7.5 mg mL^−1^, corresponding to 25% of the reaction volume, strongly reduced protein production.

Although cofactors play essential roles in metabolism, their use as CFPS supplements has been investigated only to a limited extent. We therefore evaluated *S*-adenosyl-L-methionine (SAM), biotin, thiamine pyrophosphate (TPP), vitamin B_12,_ and flavin adenine dinucleotide (FAD) added across different concentrations to the reaction. All cofactors slightly reduced GFP levels relative to the control reaction (**Figure 4E**), suggesting that additional cofactor supplementation disrupts the existing metabolic balance rather than improving CFPS under these conditions.

Motivated by these results, we performed individual titration of amino acids and NTPs using an acoustic liquid handling system (Echo 525; Beckman Coulter, Brea, CA, USA), which enables highly precise and accurate low volume transfers to enable assay miniaturization and high-throughput campaigns. Under these conditions, reaction buffers lacking individual amino acids were prepared, and each amino acid was then titrated independently. Only six amino acids (i.e., alanine, proline, serine, glutamine, tyrosine, and histidine), showed concentration-dependent effects on protein synthesis (**Figure 5A**). The remaining amino acids showed no significant effect (**Figure 5B**). This pattern is consistent with the high baseline amino acid content of non-dialyzed lysates and with amino acid interconversion during the reaction. In addition, amino acids can be synthesized from intermediates generated during maltodextrin metabolism and through interconversion reactions retained in the lysate.

**Figure 5.**
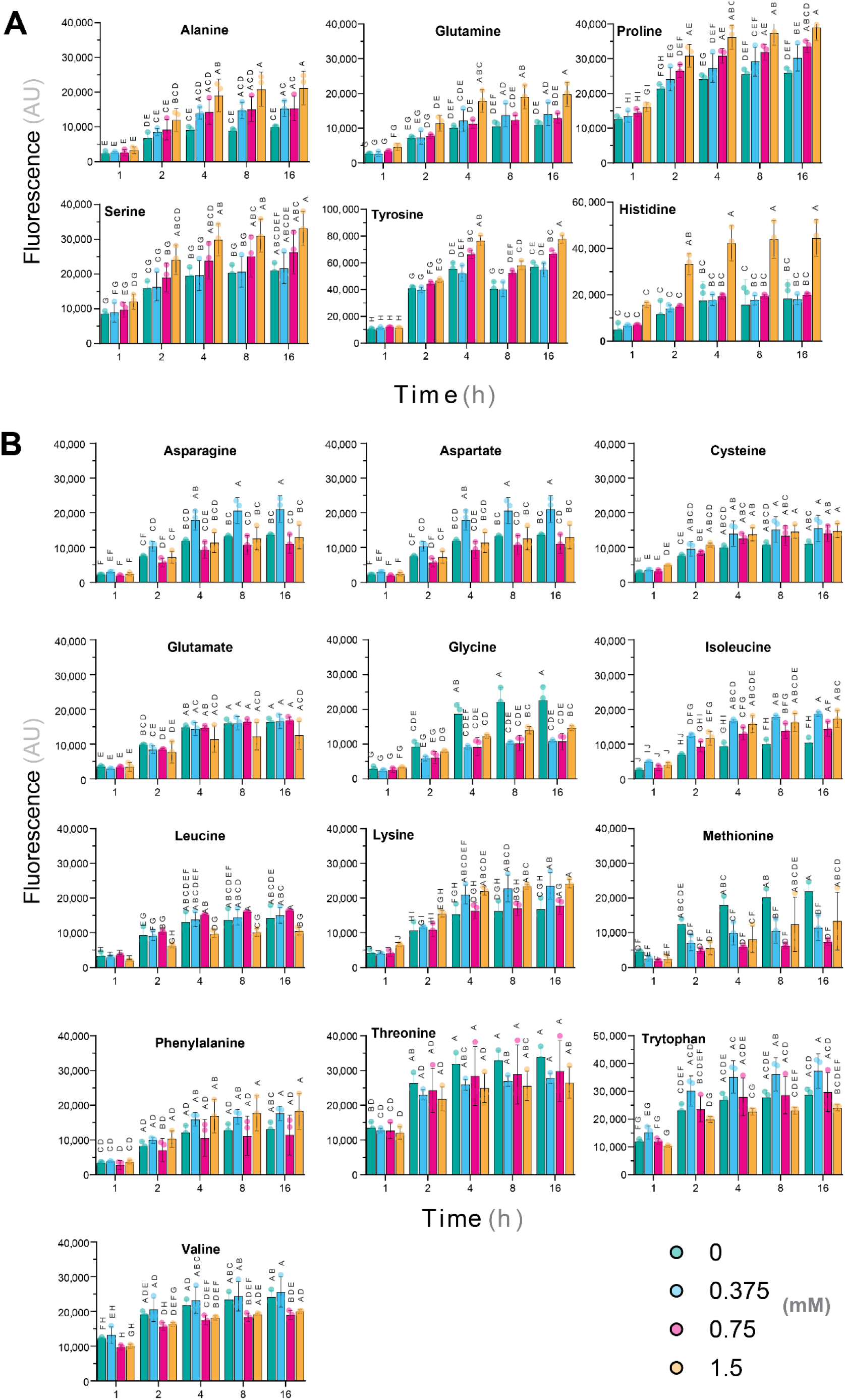
Titration of proteinogenic amino acids in CFPS. The concentration of each amino acid was varied independently using a liquid handling system, and GFP production was measured after 1, 2, 4, 8, and 16 h. A plasmid encoding deGFP was included in each reaction to monitor CFPS performance. Reactions were carried out in 384-well plates and monitored in a fluorescence plate reader. Six amino acids displayed concentration-dependent effects on GFP production **(A)**, whereas the remaining amino acids showed no consistent response **(B)**. Data represent mean values ± standard deviation from three experiments performed with independently prepared lysate batches. Technical replicates were averaged before statistical analysis. Different letters indicate statistically significant differences between groups (two-way ANOVA with Tukey’s post hoc test, *p* < 0.05; see **File S1** for details).

In contrast, all NTPs were titratable to varying degrees (**Figure 6**). Omission of any individual nucleotide strongly reduced GFP production. ATP had the largest positive effect, consistent with its central role in transcription and energy metabolism (**Figure 6A**). CTP, GTP, and UTP also positively affected protein synthesis. GTP required a minimum threshold of 1.5 mM to influence CFPS activity (**Figure 6B**). By contrast, CTP showed a strong dose-dependent response across the complete tested concentration range. Finally, CTP and UTP supplementation increased protein synthesis in a dose-dependent manner across the full tested range (**Figure 6C** and **6D**). These results identify ATP, GTP, UTP, and CTP as key regulators of reaction efficiency and reveal distinct dynamic ranges for each nucleotide.

**Figure 6.**
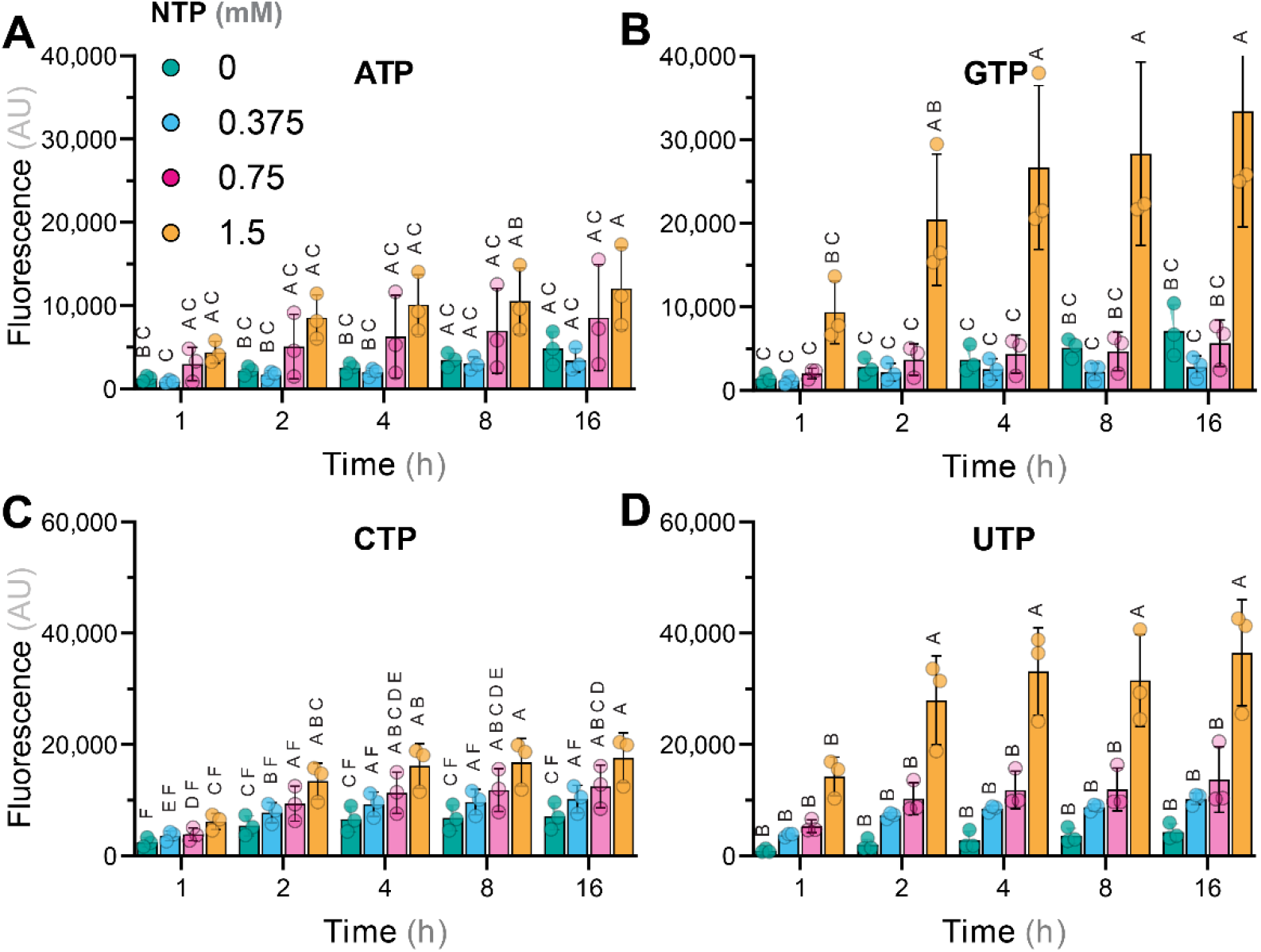
Titration of nucleoside triphosphates (NTPs) in CFPS. Individual NTPs were titrated into CFPS reactions, and deGFP production was quantified at the indicated time points. ATP **(A)**, GTP **(B)**, CTP **(C)**, and UTP **(D)** were supplemented at final concentrations of 0, 0.375, 0.75, or 1.5 mM. Data represent mean values ± standard deviation from three experiments performed with independently prepared lysate batches. Technical replicates were averaged before statistical analysis. Different letters indicate statistically significant differences between groups (two-way ANOVA with Tukey’s post hoc test, *p* < 0.05; see **File S1** for details).

### Impact of Evaporation on CFPS Performance

CFPS optimization is commonly performed in sealed microcentrifuge tubes, which provide efficient aeration while minimizing evaporation. However, these formats are poorly suited for high-throughput workflows. The headspace-to-volume ratio in microcentrifuge tubes, ca. 1:100 for a 15-μL reaction in a 1.5-mL tube, differs substantially from that of microplate formats.^3^ We therefore evaluated a 384-well plate format compatible with high-throughput CFPS applications. Each well has a total volume of 120 μL. Achieving a headspace-to-volume ratio comparable to that of microcentrifuge tubes would require a reaction volume of approximately 1.2 μL, but such low volumes are highly susceptible to evaporation and are unsuitable for extended time-course measurements. For this reason, the reaction volume was increased to 10 μL, reducing the headspace-to-volume ratio to approximately 1:12.

Three sealing approaches were tested to limit evaporation: (i) a Breathe-Easy^TM^ sealing membrane (Sigma-Aldrich Co., St. Louis, MO, USA; cat. # Z380059), which is permeable to O_2 a_nd CO_2;_ (ii) PCR sealing foil (Thermo Fisher Inc., Waltham, MA, USA; cat. # AB0558), which is impermeable to gases and liquids; and (iii) a plastic plate lid, which permits gas exchange through venting gaps. The plastic plate lid supported the highest protein production (*y*_max =_ 32,805 ± 736 AU and *K* = 2.3 ± 0.1 h; **Figure 7**). The breathable membrane and sealing foil resulted in lower protein levels. Initial production rates were similar across all conditions, indicating that oxygen availability became limiting only at later stages in sealed wells. Strongly sealed wells showed an earlier decline in protein production, with *K* = 1.4 ± 0.1 h. The plastic plate lid with venting gaps also caused substantial liquid loss, leading to variable reaction outputs. In this format, evaporation was most pronounced in the outer wells. Filling the outermost wells with water reduced evaporation and improved consistency in protein synthesis. This setup enabled stable measurements for up to 24 h at 30°C in the plate reader.

**Figure 7.**
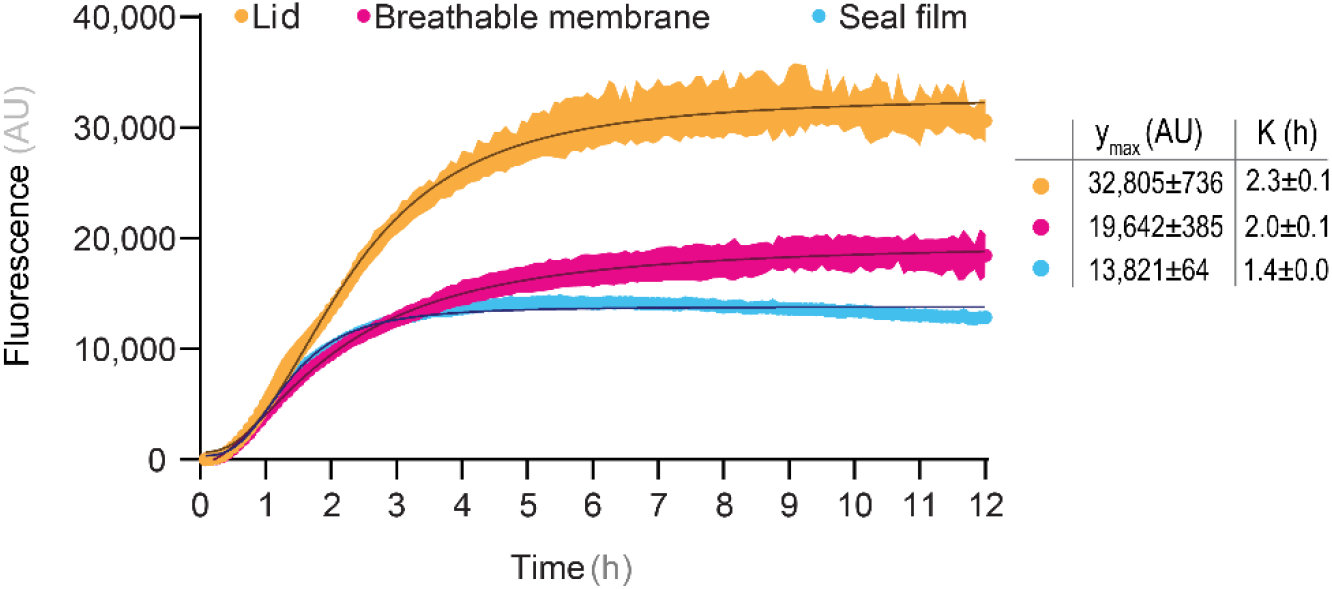
Effect of evaporation on CFPS performance. CFPS reactions were evaluated using three sealing strategies to minimize evaporation. A breathable membrane allowed gas exchange, whereas a sealing film prevented it. A plasmid encoding deGFP was included to monitor CFPS performance under each condition. Data represent mean values ± standard deviation from three experiments performed with independently prepared lysate batches. Technical replicates were averaged before statistical analysis. Detailed curve-fitting parameters are provided in **File S1**.

### Integrated Optimization of High-Throughput CFPS Reactions

The preceding experiments identified several parameters that individually improved CFPS performance in a high-throughput microplate format: 60 mM maltodextrin, 9 mg mL^−1^ lysate, 5 mM amino acids, 1.6 mM ATP and GTP, 0.9 mM UTP and CTP, and 2% (w/v) PEG. We next tested whether combining these individually optimized parameters would increase reaction output and extend productive reaction time.

Reactions containing all optimized components were assembled and monitored over time. The combined formulation produced 567 ± 64 μg mL^−1^ active deGFP (**Figure 8**). To evaluate the overall performance of the optimized CFPS, we defined the reaction completion time parameter as the time required for the fluorescence signal to reach 95% of its maximum value. Under the optimized conditions, CFPS reactions reached this threshold after ca. 14 h in 384-well plates. Fitting the protein production kinetics with **Equation 1** showed a clear improvement in *K* relative to the initial formulation (**Figure 1B**), as the optimized reaction reached half-maximal output after 2.63 h, compared with 0.88 h for the non-optimized setup.

**Figure 8.**
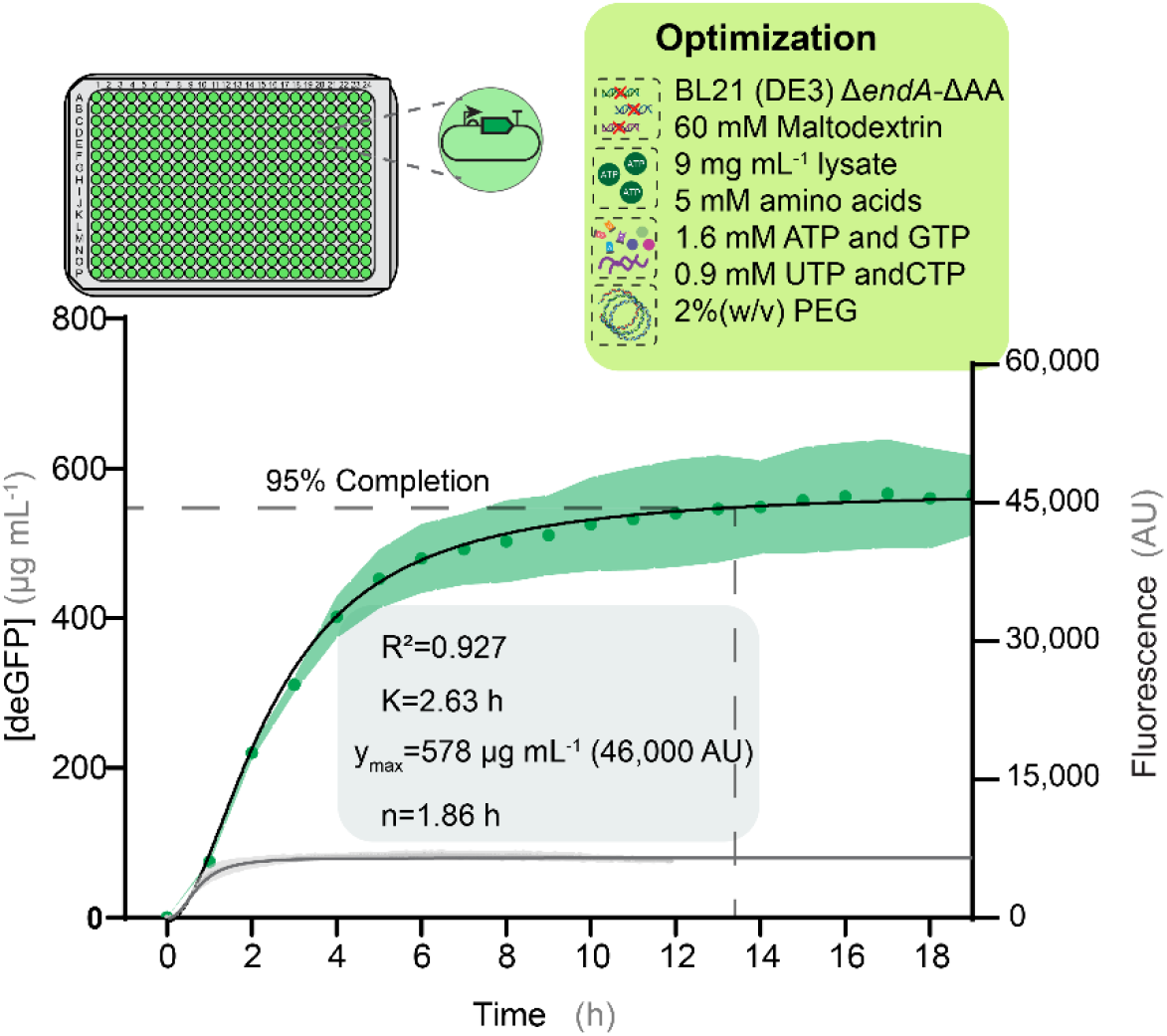
Combined parameter optimization for extended CFPS reactions. deGFP synthesis was monitored for 20 h in a 384-well plate format using the optimized reaction formulation described in the figure legend. Fluorescence was compared with an authentic commercial GFP standard to determine deGFP concentration (see **Figure S4**). Optimized reactions produced 567 ± 64 µg mL^−1^ deGFP. Reaction completion time was defined as the time required to reach 95% of the maximum fluorescence. Under optimized conditions, CFPS reactions remained active for approximately 14 h in microtiter plates. Parameters obtained by fitting the data to **Equation 1** are shown in the inset. The initial, non-optimized CFPS reaction is included in gray for comparison. Data represent mean values ± standard deviation from three experiments performed with independently prepared lysate batches. Technical replicates were averaged before statistical analysis. Detailed curve-fitting parameters are provided in **File S1**.

These results demonstrate the cumulative benefit of optimizing the three core elements of CFPS, i.e., template design, buffer composition, and lysate genotype. Each component contributed to extending the productive lifetime of the reaction, and the integrated formulation supported high protein yields in a miniaturized format compatible with high-throughput workflows.

## Conclusion

CFPS has undergone extensive development over six decades and now supports a broad range of applications.^3^ These include high-yield protein production,^66^ incorporation of non-canonical amino acids,^9^ vaccine development,^67^ and different forms of biosensing.^68^ Here, we focused on improving the suitability of CFPS for high-throughput experimentation, where predictable kinetics, extended reaction lifetimes, and compatibility with automated workflows are essential. Two applications that particularly benefit from these properties are rapid prototyping of genetic parts^4, 5^ and evaluation of biosynthetic pathways.^10, 13, 69^ Previous studies, however, have relied on large reaction volumes or microcentrifuge tubes, which limit miniaturization and reduce direct compatibility with automated systems.^70^ Droplet-based approaches can achieve small reaction volumes, but they complicate systematic optimization. By contrast, a 384-well plate format provides a practical balance between throughput, experimental accessibility, and reproducibility.

The experiments described here provide a systematic analysis of the three core components of CFPS reactions: DNA template, reaction buffer, and lysate genotype. Template stability emerged as a key determinant of performance under conditions of limited DNA availability, whereas stabilization of amino acid pools through lysate engineering contributed to extended reaction duration. Metabolomic profiling and real-time pH monitoring revealed distinct metabolic behaviors associated with different energy sources and supported the selection of maltodextrin as a robust substrate for extended reactions. Integration of these findings yielded a CFPS formulation that remained productive for approximately 14 h in a 384-well plate format while producing 567 ± 64 μg mL^−1^ active deGFP.

Collectively, these findings establish a framework for increasing the productive lifetime of CFPS in high-throughput microplate formats and provide guidance for future screening, pathway prototyping, and synthetic biology applications. Further work should evaluate the optimized system with diverse target proteins and investigate parameter interactions through multifactorial optimization to improve reaction robustness and productivity.

## Material and Methods

### Bacterial Strains, Plasmids, and Culture Conditions

All strains and plasmids used in this study are listed in **Table S1**. For cloning and genome engineering, strains were cultivated in lysogeny broth (LB) at 37°C with shaking at 200 rpm.^71^ Antibiotics were used when indicated at the following concentrations: gentamicin (Gm), 10 μg mL^−1^; kanamycin (Km), 50 μg mL^−1^; and ampicillin (Amp), 20 μg mL^−1^.

### Construction of Plasmids and Engineered Strains

All oligonucleotides used in this study are listed in **Table S2** and were purchased from Integrated DNA Technologies (IDT). Plasmids were constructed by uracil excision assembly (USER), and USER primers were designed using AMUSER software.^72^ DNA fragments were amplified with Phusion U 2X Master Mix (Thermo Scientific Inc., cat. # F533S) according to the manufacturer’s instructions. For DNA assembly, amplified fragments were mixed with USER enzyme, T4 ligase buffer, and DpnI (New England Biolabs Inc., Ipswich, MA, USA). Chemically competent *E*. *coli* DH5α λ*pir* cells were transformed with the assembled plasmids as needed. Plasmid isolation and PCR cleanup were performed using the NucleoSpin Plasmid EasyPure kit (Macherey-Nagel GmbH & Co., Düren, Germany; cat. # 740727.250) and the NucleoSpin Gel and PCR Clean-up kit (Macherey-Nagel GmbH & Co., cat. # 740609.250), respectively, according to the manufacturer’s instructions.

Several *gfp* expression plasmids were constructed from plasmid pT7 deGFP T7t.^73^ The P_T7+_ promoter was introduced by mutagenesis using the primer pair PT7+-U-F and PT7+-U-R. The T7-*_hyb10_* _t_erminator was installed using the primer pair Thyb10-U-for and Thyb10-U-rev. Plasmid pETMINI PHP-6×His was constructed by inserting a chemically synthesized fragment containing RiboJ, BCD2, and TT7*_hyb10_* _(_**Table S2**) into the pET28 vector. The insert was amplified with primers 51 and 52, while vector pET28 was amplified with primers 53 and 54. The USER-assembled plasmid was linearized again with primers 57 and 58, and PHP was amplified from pSEVA2313·PHP^49^ with primers 55 and 56 before insertion.

Functional gene disruptions in the lysate donor strain *E*. *coli* BL21 (DE3) were generated by cytosine base-editing as described by Volke et al.^55^ Briefly, sgRNA protospacers were designed using CRISPy.^74^ Single guide RNAs were inserted into vector pMBEC6 by ligation of annealed oligonucleotides into BsaI-digested vector. Multiplex guide constructs were assembled by Golden Gate cloning.^75^ *E*. *coli* DH5α λ*pir* cells were transformed with the assembled plasmids. After sequence confirmation by Sanger sequencing, the plasmids were introduced into *E*. *coli* BL21 (DE3) or its derivatives by electroporation. Colony PCR was performed to generate amplicons for Sanger sequencing and verify introduction of premature stop codons. Base-editing plasmids were cured by incubating strains harboring the plasmid in LB supplemented with 5% (w/v) sucrose. A complete list of strains constructed in this study is provided in **Table S1**.

### DNA Template Preparation for CFPS

Both plasmids and linear expression templates (LETs) were used in this study. Plasmid templates were isolated from *E*. *coli* DH5α λ*pir* after cultivation in LB for 16 h at 37°C and 200 rpm. Plasmids were purified using the NucleoSpin Plasmid EasyPure kit (Macherey-Nagel GmbH & Co.) according to the manufacturer’s instructions. DNA concentrations were quantified using a NanoDrop 2000 instrument (Thermo Scientific Inc.). Unless indicated otherwise, plasmids were used at 10 nM in each CFPS reaction.

LETs were generated by PCR using Phusion 2× High Fidelity Master Mix (Thermo Scientific Inc., cat. # F531L) with primers LET-F and LET-R (**Table S2**). These primers amplify ca. 300 bp upstream and downstream of the transcriptional unit. Amplified linear templates were purified using the NucleoSpin Gel Extraction and PCR Clean-up kit (Macherey-Nagel GmbH & Co.) according to the manufacturer’s instructions. Purified LETs were quantified using a NanoDrop 2000 instrument (Thermo Scientific Inc.).

### Cell-Free Lysate Preparation

Two methods were used for lysate preparation. Large-scale lysate was prepared by sonication using a protocol adapted from previous work.^76^ *E. coli* BL21 (DE3) or its derivatives were cultivated in 2×YT medium at 37°C and 200 rpm for 16 h. Cultures were diluted 1:100 into 100 mL of 2×YTPG medium (**Table S3**) and incubated under the same conditions. At an optical density at 600 nm (OD_600)_ of 0.6, T7 RNAP production was induced by adding isopropyl-β-D-thiogalactoside (IPTG) to a final concentration of 1 mM. Cells were grown until OD_600 r_eached ca. 3 units and harvested by centrifugation for 15 min at 4,700×*g* and 4°C. Cell pellets were washed three times with ice-cold S30 buffer freshly supplemented with 2 mM dithiothreitol (DTT) and transferred to pre-weighed 50-mL Falcon tubes. After centrifugation as above, the supernatant was removed, and wet cell pellets were weighed, flash-frozen, and subsequently stored at –80°C.

The following day, cells were thawed and resuspended in 1 mL of S30 buffer supplemented with 2 mM DTT per 1 g of wet pellet mass. Resuspended cells were aliquoted as 1.4-mL samples into 1.5-mL centrifuge tubes. Tubes were kept in an ice slurry during sonication to prevent overheating. Sonication was performed using a Vibra-Cell Sonics VCX 130 (Sonics C Materials Inc., Newtown, CT, USA) equipped with a 2-mm probe, applying 10 s *ON* and 10 s *OFF* cycles at 50% amplitude until a cumulative energy input of 800 J was reached. Sonicated samples were immediately centrifuged for 60 min at 27,000×*g* and 4°C. The supernatant was transferred to a new tube and centrifuged again for 5 min to remove residual debris. Cleared extract was distributed into PCR tubes and stored at –80°C.

Small-scale lysate was prepared by autolysis using constitutively expressed lysozyme.^77^ Lysate donor strains were transformed with plasmid pAD-LyseR and cultivated as described above, except that Amp was added. Cells were harvested at an OD_600 o_f ca. 1.5 by centrifugation for 15 min at 1,800×*g*. The supernatant was discarded, and the pellet was resuspended by vortexing in 7.5 mL of ice-cold S30 buffer supplemented with 2 mM DTT. The suspension was transferred to a pre-weighed 50-mL Falcon tube and centrifuged as above. The supernatant was removed, and the wet pellet mass was recorded. Pellets were resuspended in 2 mL of S30 buffer freshly supplemented with 2 mM DTT per 1 g of cell pellet and frozen at –80°C.

The following day, frozen cell suspensions were thawed in a room-temperature water bath. Suspensions were vortexed for 3 min and incubated in an orbital shaker for 45 min at 37°C and 600 rpm to allow lysis. This vortexing and incubation procedure was repeated once more. Cell debris was then pelleted by centrifugation at 21,000×*g* for 60 min at 4°C. The supernatant was transferred to a new tube and centrifuged again for 5 min. Cleared extract was distributed into PCR tubes and stored at –80 °C.

### Cell-Free Protein Synthesis Reactions

CFPS reagents were initially prepared as described in Didovyk *et al*.^77^ and detailed in **Tables S3**, **S5-S8**. A standard 10-μL reaction contained 4.5 μL reaction buffer, 4 μL *E*. *coli* BL21 (DE3) lysate at ca. 30 mg mL^−1^ total protein, 0.5 μL NTP mix, 0.5 μL 600 mM maltodextrin, and 0.5 μL 50 ng μL^−1^ DNA template, unless indicated otherwise. Reactions were run in 384-well plates (Corning Inc., Corning, NY, USA; cat. # 3701) at 29°C with continuous shaking for at least 16 h in a fluorescence spectrophotometer (BioTek Synergy H1 Hybrid; Agilent Technologies, Santa Clara, CA, USA). GFP fluorescence was measured every 5 min using excitation at λ_excitation =_ 485 nm and emission at λ_emission =_ 516 nm. For pH tracking with PHP, a pH-sensitive GFP variant, fluorescence was measured using excitation at λ_excitation1_ = 400 nm and λ_excitation2_ = 485 nm, with emission at λ_emission =_ 510 nm for both excitation wavelengths.

For energy source analysis, maltodextrin was replaced with alternative energy sources while maintaining the total reaction volume. To test lysates from different genetic backgrounds, three independent lysates were prepared from each strain and substituted for BL21 (DE3) lysate while keeping all other parameters constant. For lysate percentage experiments, reduced lysate volume was compensated with S30 buffer. For amino acid and PEG titration assays, reaction buffers were prepared in parallel with identical components but different amino acid or PEG concentrations, respectively. For energy source studies, maltodextrin was replaced with alternative substrates as explained in the text.

For reaction yield calculations, an 8-point standard curve was generated using the Abcam GFP Quantification Kit (Abcam Ltd., Cambridge, UK, cat. # ab235672; **Figure S4**). GFP was diluted in PBS buffer, containing 8.1 mM Na_2H_PO_4,_ 1.5 mM KH_2P_O_4,_ 137 mM NaCl, and 2.7 mM KCl at pH = 7.5. Fluorescence was measured in 384-well plates using a BioTek Synergy H1 instrument with λ_excitation =_ 485 nm and λ_emission =_ 528 nm.

### Metabolomic Analyses

Metabolomics was performed for selected energy sources and engineered strains following the procedure described by Rasor et al.^78^ Four 10-µL reactions were prepared in a 384-well microtiter plate and incubated for 4 h. At 0 h, 5 µL were collected from each reaction and pooled; the remaining reaction material was pooled after 4 h. Reactions were immediately quenched by adding 100 μL of quenching solution containing 10 mM tributylamine and 10 mM acetic acid in 5% (v/v) methanol, 2% (v/v) isopropanol, and 93% (v/v) water. Samples were centrifuged at 13,000×*g* for 10 min to precipitate insoluble material, and the supernatant was transferred to a new reaction tube. Samples were placed in an Eppendorf Concentrator Plus (Sartorius AG, Göttingen, Germany) and solvents were evaporated at room temperature for 90 min. After complete drying, 80 µL of deionized water was added to each sample. Samples were frozen and stored at –80°C until further processing. Samples were transferred to injection tubes and analyzed by liquid chromatography-mass spectrometry (LC-MS) as reported previously.^79^ Chromatograms were analyzed using MultiQuant software (AB Sciex LLC, Marlborough, MA, USA), and Prism 9.5 software (GraphPad Software Inc., San Diego, CA, USA) was used to plot results for selected metabolites.

### Real-Time pH Tracking of CFPS Reactions

The pH-sensitive GFP variant pHluorin2 was used to track real-time changes of the pH in CFPS reactions.^49^ *E. coli* BL21 (DE3) was transformed with plasmid pETMINI-PHP-6×His, which carries a codon-optimized pHluorin2 gene. Cells harboring the plasmid were grown in LB supplemented with Km at 37°C. At OD_600 =_ 0.6, expression of the T7 RNAP gene was induced, enabling production of His-tagged PHP. Cells were cultivated for an additional 6 h at 37°C and harvested by centrifugation at 4,500×*g* for 15 min. The supernatant was discarded, and the cell pellet was resuspended in buffer A, containing 300 mM NaCl, 20 mM imidazole, and 10 mM HEPES at pH = 7.5. The sample was transferred to a 50-mL Falcon tube, placed on ice, and sonicated using a Vibra-Cell Sonics VCX 130 equipped with a 6-mm probe, applying 10 s *ON* and 10 s *OFF* cycles for 2 min. After sonication, the sample was clarified by centrifugation at 27,000×*g* for 1 h at 4°C, and the supernatant was transferred to a new 50-mL Falcon tube. For protein purification, 1 mL of HisPur cobalt resin (Thermo Scientific Inc., cat. # 89964) was equilibrated with 5 mL of buffer A and incubated for 3 min to allow resin sedimentation. The liquid above the resin was carefully removed, and the protein sample was added to the resin. The mixture was rotated for 1 h at 4°C to allow protein binding; the non-binding protein fraction was removed by washing twice with 10 mL of buffer A. Bound protein was eluted twice with 2 mL of buffer B, containing 300 mM NaCl, 500 mM imidazole, and 10 mM HEPES at pH = 7.5. The buffer was exchanged to buffer C, containing 50 mM sodium phosphate at pH = 7.8, using a 10-kDa Amicon Ultra centrifugal filter.

To calibrate the pH response, the fluorescence ratio of purified 2.5 μM pHluorin2 between the two excitation maxima, 400 nm and 485 nm, with emission at 510 nm, was measured in S30 buffer adjusted to different pH values (**Figure S1**). To avoid crosstalk between PHP and GFP in CFPS reactions, a non-fluorescent GFP mutant, dGFP, was used in samples for pH measurement. In all experiments, purified pHluorin2 was added to each CFPS reaction at a final concentration of 2.5 μM.

### Data Representation and Statistical Analysis

Experimental CFPS data were fitted using the following equation:

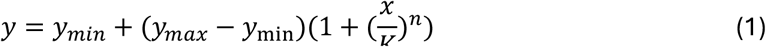

where *y* is the CFPS output in AU or μg mL^−1^, *y_min_* is the background fluorescence output, *y_max_* is the maximum fluorescence output, *x* is the reaction time, *K* is the time to reach half of the maximum output and *n* is the cooperativity index.

Figures were prepared using GraphPad Prism 9.5, Illustrator and Inkscape. Statistical analyses were performed using GraphPad Prism analysis tools. Details of statistical analysis are presented in **File S1**. Statistical significance was interpreted according to the following *p*-value thresholds: *p* < 0.0001, ****; *p* < 0.001, ***; *p* < 0.01, **; *p* < 0.05, *; and *p* > 0.05, non-significant. For experiments including large data sets (amino acids and nucleotide titration) significant differences are given in compact letter format, with *p* < 0.05. All experiments were performed with independent biological replicates, and data are presented as mean ± standard deviation from independent experiments.

## AUTHORS’ CONTRIBUTIONS

**EUB:** Investigation, Data curation, Methodology, Validation, Visualization, Writing – original draft; **BZ:** Investigation; **PIN:** Conceptualization, Resources, Funding acquisition, Supervision, Project administration, Writing – review & editing; **DCV:** Conceptualization, Investigation, Data curation, Methodology, Validation, Visualization, Supervision, Writing – original draft & editing.

## ACKNOWLEDGMENTS

The work was supported by The Novo Nordisk Foundation through grants NNF20CC0035580, *LiFe* (NNF18OC0034818), *TARGET* (NNF21OC0067666), FM·*Pseudomonas* (NNF24OC0061501), *NovoF* (NNF23OC0083631), SCOUT (NNF25OC0103413) and NNF24SA0100980, and the

European Union’s Horizon 2020 Research and Innovation Programme under grant agreement No. 101082046 (*TOLERATE*).

## DATA AVAILABILITY

The data, strains, and plasmids that support the findings of this study are available from the corresponding authors upon reasonable request.

## ETHICS STATEMENT

The work presented in this article follows all prevailing local, national, and international regulations and conventions, and normal scientific ethical practices.

