## Supplementary Material for "Multiparameter optimization extends the lifetime of cell-free protein synthesis in a high-throughput format"

by

Eray U. Bozkurt, Baptiste Zanchet, Pablo I. Nikel* and Daniel C. Volke*

### SUPPLEMENTARY TABLES AND FIGURES

**Table S1.** Bacterial strains and plasmids used in this study.

| Name | Relevant characteristics[a] | Reference or source |
|---|---|---|
| | **Strains** | |
| ***E. coli* DH5a λpir** | Cloning host; $F^{-}$ $l^{-}$ endA1 glnX44(AS) thiE1 recA1 relA1 spoT1 gyrA96($Nal^{R}$) rfbC1 deoR nupG F80(lacZDM15) D(argF-lac)U169 hdR17($r_K^{-}$ $m_K^{+}$), λpir lysogen | [1] |
| ***E. coli* BL21 (DE3)** | Protein production and lysate preparation host; F– *ompT* $hsdS_B$ ($r_B$–, $m_B$–) *gal dcm* (DE3) | [2] |
| ***E. coli* BL21 (DE3) Δ*csdA*** | Derivate of *E. coli* BL21 (DE3) with premature stop codons inserted into *csdA* | This study |
| ***E. coli* BL21 (DE3) Δ*endA*** | Derivate of *E. coli* BL21 (DE3) with premature stop codons inserted into *endA* | This study |
| ***E. coli* BL21 (DE3) Δ*mazF*** | Derivate of *E. coli* BL21 (DE3) with premature stop codons inserted into *mazF* | This study |
| ***E. coli* BL21 (DE3) Δ*gorA*** | Derivate of *E. coli* BL21 (DE3) with premature stop codons inserted into *gorA* | This study |
| ***E. coli* BL21 (DE3) ΔAA** | Derivate of *E. coli* BL21 (DE3) with premature stop codons inserted into *gshA tnaA speA sdaA sdaB* | This study |
| ***E. coli* BL21 (DE3) Δ*rnE*** | Derivate of *E. coli* BL21 (DE3) with premature stop codons inserted into *rnE* | This study |
| ***E. coli* BL21 (DE3) Δ*rnB*** | Derivate of *E. coli* BL21 (DE3) with premature stop codons inserted into *rnB* | This study |
| ***E. coli* BL21 (DE3) Δ*csdA* Δ*endA*** | Derivate of *E. coli* BL21 (DE3) with premature stop codons inserted into *csdA and endA* | This study |
| ***E. coli* BL21 (DE3) Δ*endA* Δ*rnB*** | Derivate of *E. coli* BL21 (DE3) with premature stop codons inserted into *endA and rnB* | This study |

| | | |
|---|---|---|
| ***E. coli* BL21 (DE3) Δ*csdA* Δ*rnB*** | Derivate of *E. coli* BL21 (DE3) with premature stop codons inserted into *csdA and rnB* | This study |
| ***E. coli* BL21 (DE3) Δ*csdA* ΔAA** | Derivate of *E. coli* BL21 (DE3) with premature stop codons inserted into *csdA gshA tnaA speA sdaA sdaB* | This study |
| ***E. coli* BL21 (DE3) Δ*endA* ΔAA** | Derivate of *E. coli* BL21 (DE3) with premature stop codons inserted into *endA gshA tnaA speA sdaA sdaB* | This study |
| ***E. coli* BL21 (DE3) Δ*gorA* ΔAA** | Derivate of *E. coli* BL21 (DE3) with premature stop codons inserted into *gorA gshA tnaA speA sdaA sdaB* | This study |
| **Plasmids** | | |
| **pT7·msfGFP-$T_0$** | *msfGFP* expression plasmid; $P_{T7}$->*msfGFP*-$T_{To}$, *oriV(pBBR1),* $Km^R$, (plasmid called pSEVA2325M after SEVA nomenclature) | [3] |
| **pT7·GFPmut3-$T_{T7}$** | *GFPmut3* expression plasmid; $P_{T7}$->*GFPmut3*-$T_{T7}$, *oriV(pBR322),* $Amp^R$ | This study |
| **pT7·deGFP-$T_{T7}$** | *deGFP* expression plasmid; $P_{T7}$->*deGFP*-$T_{T7}$, *oriV(pBR322),* $Amp^R$ | Gift from Richard Murray (Addgene plasmid # 67741) |
| **pT7+·deGFP-$T_{T7}$** | *deGFP* expression plasmid; $P_{T7+}$->*deGFP*-$T_{T7}$, *oriV(pBR322),* $Amp^R$ | This study |
| **pT7·deGFP-$T_{T7hyb10}$** | *deGFP* expression plasmid; $P_{T7}$->*deGFP*-$T_{T7-hyb}$, *oriV(pBR322),* $Amp^R$ | This study |
| **pT7+·deGFP- $T_{7hyb10}$** | *deGFP* expression plasmid; $P_{T7+}$->*deGFP*-$T_{T7-hyb}$, *oriV(pBR322),* $Amp^R$ | This study |
| **pS23T7·dGFP** | *Non-fluorescent msfgfp* (msfGFP$^{T65G, S73A, M79I, Q81R, M88L, N149K, K206V, S208I, M219A, M233K}$) *expression plasmid;* $P_{T7}$->*dGFP*-$T_{T7,}$ *oriV(pBBR1),* $Kan^R$ | This study |
| **PHP** | $P_{EM7}$-> pHIorin2, *oriV(RSF1010),* $Km^R$, (plasmid called pSEVA2313·PHP after SEVA nomenclature) | [4] |
| **pETMINI · PHP-6xHis** | $P_{T7}$->RiboJ-BCD2-PHP-6xHis-$T_{T7-hyb,}$ *oriV(pBR322),* $Km^R$ | This study |

| | | |
|---|---|---|
| **pAD-LyseR** | *Autolysis plasmid,* $P_{bla}$-> *endolysine* (phage lambda, gene R), *oriV(pBR322),* $\text{Amp}^{R}$ | [5] |
| **pMBEC6** | Multiplex cytosine base-editing vector bearing the uracil glycosylase inhibitor *(ugi)* gene, the monomeric superfolder green fluorescent protein *(msfGFP)* gene, and the *Cas6* endoribonuclease gene*; oriV(pRO1600/ColE1);* $\text{Gm}^{R}$ | [6] |
| **pMBEC6·*csdA*** | Derivative of vector pMBEC6 bearing a gRNA with *csdA* spacers (Q131*, Q132*); $\text{Gm}^{R}$ | This study |
| **pMBEC6·*endA*** | Derivative of vector pMBEC6 bearing a gRNA with *endA* spacers (W82*); $\text{Gm}^{R}$ | This study |
| **pMBEC6·*mazF*** | Derivative of vector pMBEC6 bearing a gRNA with *mazF* spacers *(*R4*); $\text{Gm}^{R}$ | This study |
| **pMBEC6·*gorA*** | Derivative of vector pMBEC6 bearing a gRNA with *gorA* spacers (W54*); $\text{Gm}^{R}$ | This study |
| **pMBEC6·*rnE*** | Derivative of vector pMBEC6 bearing a gRNA with *rnE* spacers (W191*); $\text{Gm}^{R}$ | This study |
| **pMBEC6·*rnB*** | Derivative of vector pMBEC6 bearing a multiplex gRNA with *rnB* spacers (Q149*, W234*); $\text{Gm}^{R}$ | This study |
| **pMBEC6·AA** | Derivative of vector pMBEC6 bearing a multiplex gRNA with *gshA (W56*), tnaA (Q203*), speA (Q305*), sdaA (Q100*), sdaB (Q88*)* spacers; $\text{Gm}^{R}$ | This study |

**Table S2**. Oligonucleotides and synthetic DNA fragments used in this study.

| Oligo # | Oligo name | Sequence (5'→3') | Purpose |
|---|---|---|---|
| **1** | LET-for | GACGTTGTAAAACGACGGCCAG | Amplification of linear expression template |
| **2** | LET-rev | AGCGGATAACAATTTCACACAGGA | |
| **3** | PT7+-U-for | ATTTCCCUATAGTGAGTCGTATTAATTTCGCG | pT7+ promoter cloning |
| **4** | PT7+-U-for | AGGGAAAUAACAACGGTTTCCCTCTAGAA | |
| **5** | Thyb10-U-for | ATCTGTUGAAAAAAAACAGATAACAGATACCGAAG TATCTGTTATCTTTATTGCTCAGCGGTGGC | Thyb10 terminator cloning |
| **6** | Thyb10-U-rev | AACAGAUAGCCGCGTTCGCGCGGCTATCTGTTTT TTTTCTGAAAGGAGGAACTATATCCG | |
| **7** | endA_gRNA-for | GTGGGTGTTCCCACTCTACGCGGC | Insertion of spacer for *endA* editing into pMBEC6 |
| **8** | endA_gRNA-rev | AAACGCCGCGTAGAGTGGGAACAC | |
| **9** | csdA_gRNA-for | GTGGCCAACAAACTGGAGCCAAAG | Insertion of spacer for csdA editing into pMBEC6 |
| **10** | csdA_gRNA-rev | AAACCTTTGGCTCCAGTTTGTTGG | |
| **11** | gshA-pos1-gg-for | ATCGAGGTCTCCGTGGAATCCATTTGTGCGTCAGT GGTTTTAGAGCTAGAAATAGC | Insertion of spacer for *gshA* editing into pMBEC6 |
| **12** | tnaA-pos2-gg-for | ATCGAGGTCTCCAGGTAGGTGGTCAGCCGGTTTC ACGTTTTAGAGCTAGAAATAGC | Insertion of spacer for *tnaA* editing into pMBEC6 |
| **13** | speA-pos3-gg-for | ATCGAGGTCTCCGCGGAATATTCAGTGCTTCGAC GTGTTTTAGAGCTAGAAATAGC | Insertion of spacer for *speA* editing into pMBEC6 |

| | | | |
|---|---|---|---|
| **14** | sdaA-pos4-gg-for | ATCGAGGTCTCCTTGTGGCACAGGGACGGCATGAAGGTTTTAGAGCTAGAAATAGC | Insertion of spacer for *sdaA* editing into pMBEC6 |
| **15** | sdaB-pos5-gg-for | ATCGAGGTCTCCTGTTATTCAGGATGTGAATACTCAGTTTTAGAGCTAGAAATAGC | Insertion of spacer for *sdaB* editing into pMBEC6 |
| **16** | gorA_gRNA-for | GTGGGCGTGCCACATCACTTTTTT | Insertion of spacer for *gorA* editing into pMBEC6 |
| **17** | gorA_gRNA-rev | AAACAAAAAAGTGATGTGGCACGCAAAC | |
| **18** | mazF_gRNA-for | GTGGAAGCCGATACGTACCCGATA | Insertion of spacer for *gorA* editing into pMBEC6 |
| **19** | mazF_gRNA-rev | AAACTATCGGGTACGTATCGGCTTAAAC | |
| **20** | rne_gRNA-for | GTGGTGGCTTCCCAGTGTTTCAGA | Insertion of spacer for *rnE* editing into pMBEC6 |
| **21** | rne_gRNA-rev | AAACTCTGAAACACTGGGAAGCCA | |
| **22** | rnb1-pos1-gg-for | ATCGAGGTCTCCGTGGCTGACACAATACATCACTTTGTTTTAGAGCTAGAAATAGC | Insertion of first spacer for *rnE* editing into pMBEC6 |
| **23** | rnb2-pos2-gg-for | ATCGAGGTCTCCAGGTAGCAATCCACGCGGTTGGATGTTTTAGAGCTAGAAATAGC | Insertion of second spacer for *rnE* editing into pMBEC6 |
| **24** | gRNA-Position1-gg-rev | ATCGAGGTCTCCACCTTTAGCTGCCTATACGGCAGT | Standard Golden Gate oligonucleotides for assembly of multipe gRNAs in pMBEC |
| **25** | gRNA-Position2-gg-rev | ATCGAGGTCTCCCCGCTTAGCTGCCTATACGGCAGT | |
| **26** | gRNA-Position3-gg-rev | ATCGAGGTCTCCACAATTAGCTGCCTATACGGCAGT | |

| | | | |
|---|---|---|---|
| **27** | gRNA-Position4-gg-rev | ATCGAGGTCTCCAACATTAGCTGCCTATACGGCAGT | |
| **28** | gRNA-lastPosition-gg-rev | ATCGAGGTCTCCAAACTTTCTTAGCTGCCTATACGG | |
| **29** | endA-seq-for | CAGCATTTTCCGGCCCGGCG | Sequencing *endA* |
| **30** | endA-seq-rev | GCTGGTGGTTCGGCAGCTTT | |
| **31** | csdA_seq-for | GCCTGACCGCGCGTTATGAA | Sequencing *csdA* |
| **32** | csdA_seq-rev | CGCCAGATCCGGGCAACC | |
| **33** | gshA_seq-for | TTTGCATCAGCGCGCCGTAG | Sequencing *gshA* |
| **34** | gshA_seq-rev | TACAGCGTGGGCTGGAGCGC | |
| **35** | tnaA_seq-for | GCCTTCGATACGGGCGTGCG | Sequencing *tnaA* |
| **36** | tnaA_seq-rev | TGGCGGACATCGCCAGCATA | |
| **37** | speA_seq-for | ACCGTCGGATGCGGCAGACC | Sequencing *speA* |
| **38** | speA_seq-rev | GCACGTCTGGCTTCGCAGGG | |
| **39** | sdaA_seq-for | GGGATTGGTCCCTCATCTTCCC | Sequencing *sdaA* |
| **40** | sdaA_seq-rev | TTTGAACGGATACGGCACGC | |
| **41** | sdaB_seq-for | GGCCACCACACTGATATCGCC | Sequencing *sdaB* |
| **42** | sdaB_seq-rev | GGCCAGAGAGTGACAGCCCG | |

| | | | |
|---|---|---|---|
| **43** | gorA_seq-for | ATTACATCGCCATCGGCGGC | Sequencing *gorA* |
| **44** | gorA_seq-rev | GCCCGGAATATCCGGGTGGC | |
| **45** | rne_seq-for | GGAAGGCGGACTCGATCTGTG | Sequencing *rnE* |
| **46** | rne_seq-rev | TCTCGCCGTATCGAAGGCGA | |
| **47** | mazF_seq-for | GCCATCACGTTCCTGACCGG | Sequencing *mazF* |
| **48** | mazF_seq-rev | GGGGAAATTCACCGGCGGTG | |
| **49** | rnb-seq-for | CGCCTTAGCGAAAAGGGCGTC | Sequencing *rnB* |
| **50** | rnb-seq-rev | CGTTCCTGACTCGTTTCGTGGG | |
| **51** | T7-riboJ-TT7hyb10-for | AGTTTTTCUAAAAAAAAACAGATAGCCG | Assembly of pETMini |
| **52** | T7-riboJ-TT7hyb10-rev | AAACTAAUACGACTCACTATAGGGAT | |
| **53** | pETmini-bb-U-for | ATTAGTTUTTCCATAGGCTCCGCCCC | |
| **54** | pETmini-bb-U-rev | AGAAAAACUCATCGAGCATCAAATGAA | |
| **55** | PHP-U-for | AGCAAAGGUGAAGAATTGTTCACCGGTGTG | Assembly of pETMINI PHP-6xHis |
| **56** | PHP-U-rev | ATCTTTAAUGGTGGTGGTGGTGGTGGG | |
| **57** | pETmini-U-for | ATTAAAGAUAACAGATACTTCGGTATC | |
| **58** | pETmini-U-rev | ACCTTTGCUCATTAGAAAACCTCCTTA | |

| DNA fragment | | | |
|---|---|---|---|
| **59** | T7-riboJ-TT7hyb10 | TAATACGACTCACTATAGGGATAATAGCTGTCACCGGATGTGCTTTCCGGTCTGATGAGTCCGTGAGGACGAAACAGCCTCTACAAATAATTTTGTTTAAGGGCCCAAGTTCACTTAAAAAGGAGATCAACAATGAAAGCAATTTTCGTACTGAAACATCTTAATCATGCTAAGGAGGTTTTCTAATG | T7Pr (black)-RiboJ (orange)-$T_{T7hyb10}$ (blue) insert for pETMini |

**Table S3**. Recipe for buffers used in this study.

| **Buffers/ Reagents** | **Recipe** | **Preparation** |
|---|---|---|
| S30 Buffer | 10 mM (1.81 g $L^{-1}$) Tris(hydroxymethyl)aminomethane (Tris) - acetate pH 8.2, 14 mM (3 g $L^{-1}$) $Mg(CH_3COO)_2$, 60 mM (5.89 g $L^{-1}$) K $CH_3COO$, 1 M dithiothreitol (DTT) | pH 8.2. Filter-sterilized. DTT is added prior to use to 2 mM final concentration |
| 2YPTG media | 5 g $L^{-1}$ NaCl, 16 g $L^{-1}$ tryptone, 10 g $L^{-1}$ yeast extract, 7 g $L^{-1}$ $K_2HPO_4$, 3 g $L^{-1}$ $KH_2PO_4$, 100 mM (18 g $L^{-1}$) glucose | Combine all ingredients except glucose and add water to 90% of final volume. Adjust pH to 7.2 with 5 N NaOH and autoclave. Add 100 mL of 1 M glucose after autoclaving |
| LB | 10 g $L^{-1}$ tryptone, 5 g $L^{-1}$ yeast extract, and 10 g $L^{-1}$ NaCl | Sterilize by autoclaving |

**Table S4**. A detailed explanation of the modifications of *E. coli* BL21 (DE3) strains.

| Gene | Details |
|---|---|
| ***endA*** | Endonuclease-1—endonucleolytic cleavage to 5`-phosphodinucleotide and 5´-phosphooligonucleotide end-products |
| ***csdA*** | RNA helicase CsdA—mRNA decay initiation |
| ***mazF*** | Endonuclease MazF- Endoribonucleolytic cleavage of mRNA |
| ***gorA*** | Glutathione reductase—reduction of glutathione disulfide to glutathione |
| ***gshA*** | Glutamate-cysteine ligase |
| ***tnaA*** | Hydrolytic β-elimination of L-tryptophan |
| ***speA*** | Arginine decarboxylase--- biosynthesis of agmatine from arginine |
| ***sdaA- sdaB*** | L-serine dehydratase 1 and 2 – deamination of L-serine and L-threonine |

**Table S5**. General CFPS reaction setup. The reactions were adjusted accordingly when testing an element of CFPS. For example, in energy source studies, Maltodextrin was replaced by related energy source.

| CFPS Setup | | |
|---|---|---|
| **Brand** | **Reagent** | **Volume(µL)** |
| - | Reaction Buffer | 4.5 |
| - | Lysate | 4 |
| Sigma (419680) | 600 mM Maltodextrin | 0.5 |
| GE(GE27-2025-01) | NTPs (32 mM ATP&GTP; 18 mM UTP&CTP ) | 0.5 |
| - | DNA | 0.5 |

**Table S6**. Preparation of reaction buffer used in CFPS reactions.

| Reaction Buffer | | |
|---|---|---|
| **Volume (ul)** | **Stock concentration (x)** | **Reagent** |
| 714.3 | 14 | Solution I |
| 2500 | 4 | 4X amino acid mix |
| 500 | 20 | 40% w/v PEG-8000 |
| 303 | 33 | 2 M K-glutamate |
| 25 | 400 | 1 M Mg-glutamate |
| 100 | 100 | 100 mM DTT |

**Table S7.** Preparation of 4X amino acid mix used in reaction buffer formulation.

| 4X Amino acid mix | | | | | |
|---|---|---|---|---|---|
| **Amino acid** | **Abbreviation** | | **MW (g mol$^{-1}$)** | **Final concn. (mM)** | **Amount to add (mg)** |
| Alanine | Ala | A | 89.1 | 24 | 85.536 |
| Arginine | Arg | R | 174.2 | 24 | 167.232 |
| Asparagine | Asn | N | 132.1 | 24 | 126.816 |
| Aspartate | Asp | D | 133.1 | 24 | 127.776 |
| Cysteine | Cys | C | 121.2 | 24 | 116.352 |
| Glutamate | Glu | E | 147.1 | 24 | 141.216 |
| Glutamine | Gln | Q | 146.2 | 24 | 140.352 |
| Glycine | Gly | G | 75.1 | 24 | 72.096 |
| Histidine | His | H | 155.2 | 24 | 148.992 |
| Isoleucine | Ile | I | 131.2 | 24 | 125.952 |
| Leucine | Leu | L | 131.2 | 20 | 104.96 |
| Lysine | Lys | K | 146.2 | 24 | 140.352 |
| Methionine | Met | M | 149.2 | 24 | 143.232 |
| Phenylalanine | Phe | F | 165.2 | 24 | 158.592 |
| Proline | Pro | P | 115.1 | 24 | 110.496 |
| Serine | Ser | S | 105.1 | 24 | 100.896 |
| Threonine | Thr | T | 119.1 | 24 | 114.336 |
| Tryptophan | Trp | W | 204.2 | 24 | 196.032 |
| Tyrosine | Tyr | Y | 181.2 | 24 | 173.952 |
| Valine | Val | V | 117.1 | 24 | 112.416 |
| | | | volume to dissolve (mL): | | 40 |
| **All amino acids were purchased from Sigma Aldrich. Final pH 6.8. Aliquoted and stored at -80 °C.** | | | | | |

**Table S8.** Preparation of 14X solution used in reaction buffer formulation.

| **14x Solution I** | | | | |
|---|---|---|---|---|
| Concentration | Units | Name | MW (g mol$^{-1}$) | added weight (mg) |
| 700 | mM | HEPES (free acid) | 238.3 | 667.20 |
| 2.8 | mg mL$^{-1}$ | tRNA (Roche MRE600, 10109541001) | | 11.20 |
| 3.64 | mM | CoA (Sigma C4282) | 767.53 | 11.20 |
| 4.62 | mM | NAD (Sigma N6522) | 663.43 | 12.30 |
| 10.5 | mM | cAMP (Sigma A9501) | 329.21 | 13.80 |
| 0.95 | mM | Folinic acid (Sigma F7878) | 511.5 | 1.90 |
| 14 | mM | Spermidine (Sigma 85558) | 145.25 | 8.10 |
| HEPES is resuspended in 2 mL deionized water and pH is arranged to 8. Subsequently, all other compounds were added and water is added to a final volume of 4 mL. The pH was adjusted to 7.6 and the solution was filter-sterilized, aliquoted to 50 µL and stored at -80°C until use. | | | | |

**Table S9.** Full name of detectable compounds in metabolomics analysis.

| Abbreviations of chemicals | |
|---|---|
| **Compound name** | **BIGG ID** |
| 2,3-Bisphosphoglyceric acid | 23dpg |
| 2-methylcitrate | 2mcit |
| 2-Oxobutanoate | 2obut |
| 2-Oxoglutarate | akg |
| 3',5'-Cyclic GMP | 35cgmp |
| 5-Amino-1-(5-Phospho-ribosyl)imidazole-4-carboxamide | AICAr |
| 5-Oxoproline | 5oxpro |
| 5-Phospho-alpha-ribose 1-diphosphate | prpp |
| 6-Phospho-gluconate | 6pgc |
| Acetyl phosphate | actp |
| Acetyl-Coenzyme A | accoa |
| Adenine | ade |
| Adenosine | adn |
| ADP | adp |
| ADPglucose | adpglc |
| Alanine | ala-L |
| AMP | amp |
| Arginine | arg-L |
| Asparagine | asn-L |
| Aspartic acid | asp-L |
| ATP | atp |
| Biotin | btn |
| cAMP | camp |
| Chorismate | chor |
| cis-Aconitate | acon-c |

| | |
|---|---|
| Citrate | cit |
| Citrulline | citr-L |
| CMP | cmp |
| Coenzyme A | coa |
| CTP | ctp |
| Cystine | Lcystin |
| Cytidine | cytd |
| dADP | dadp |
| dAMP | damp |
| dATP | datp |
| dCDP | dcdp |
| dCMP | dcmp |
| dCTP | dctp |
| dGDP | dgdp |
| dGMP | dgmp |
| DGTP | dgtp |
| Dihydroxyacetone phosphate | dhap |
| dIMP | dimp |
| DITP | ditp |
| dTDPglucose | dtdpglu |
| dUMP | dump |
| dUTP | dutp |
| Flavin adenine dinucleotide oxidized | fad |
| Folate | fol |
| Fructose 1,6-bisphosphate | fdp |
| Fructose 1-phosphate | f1p |
| Fructose 6-phosphate | f6p |
| Fumarate | fum |
| GDP | gdp |
| Glucosamine 6-phosphate | gam6p |

| | |
|---|---|
| Glucose 1-phosphate | g1p |
| Glucose 6-phosphate | g6p |
| Glutaconate | glutacon |
| Glutamate | glu-L |
| Glutamine | gln-L |
| Glyceraldehyde-3-phosphate | g3p |
| Glycerol 3-phosphate | glyc3p |
| Glycolate | glyclt |
| Glyoxylate | glx |
| GMP | gmp |
| GTP | gtp |
| Guanine | gua |
| Guanosine | gsn |
| Histidine | his-L |
| Hypoxanthine | hxan |
| IMP | imp |
| Inosine | ins |
| Isocitrate | icit |
| ITP | itp |
| Lactate | lac-L |
| Malate | mal-L |
| Malonyl-Coenzyme A | malcoa |
| Methionine | met-L |
| Methylmalonate | mmal |
| Nicotinamide adenine dinucleotide | nad |
| Nicotinamide adenine dinucleotide - reduced | nadh |
| Nicotinamide adenine dinucleotide phosphate | nadp |
| Nicotinamide adenine dinucleotide phosphate - reduced | nadph |
| Ornithine | orn |

| | |
|---|---|
| Oxalate | oxa |
| Oxaloacetate | oaa |
| Oxidized glutathione | gthox |
| Phenylalanine | phe-L |
| Phenylpyruvate | phpyr |
| Phosphoenolpyruvate | pep |
| Pool 2 phosphoglycerate/3-phosphoglycerate | Pool_2pg_3pg |
| Pool Citrate/Isocitrate | Pool_cit_icit |
| Pool fructose/glucose | Hexose_Pool_fru_glc-D |
| Pyruvate | pyr |
| Reduced glutathione | gthrd |
| Riboflavin | ribflv |
| Ribose 5-phosphate | r5p |
| Ribulose 5-phosphate | ru5p-D |
| Sedoheptulose 7-phosphate | s7p |
| Serine | ser-L |
| Shikimate | skm |
| Succinate | succ |
| Succinyl-Coenzyme A | succoa |
| Threonine | thr-L |
| Thymine | thym |
| TMP | dtmp |
| Tryptophan | trp-L |
| TTP | dttp |
| Tyrosine | tyr-L |
| UDP | udp |
| UDP-D-glucuronate | udpglcur |
| UDPglucose | udpg |
| UMP | ump |
| Uracil | ura |

| | |
|---|---|
| Urate | urate |
| Uridine | uri |
| UTP | utp |
| Xanthine | xan |

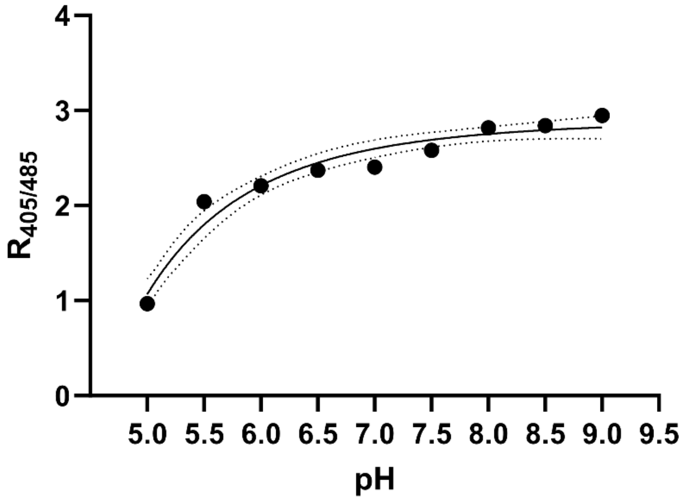


**Figure S1 |** Calibration curve of pH sensor PHP. The protein was purified and diluted in S30 buffer at different pH. Fluorescence was recorded at 405 nm and 485 nm excitation, and the ratio of both signals was calculated.

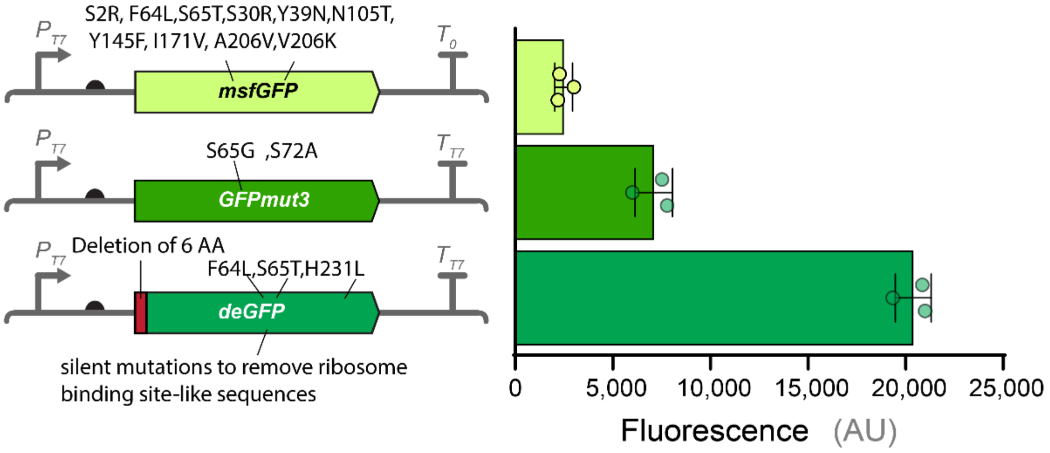


**Figure S2** | Constructs encoding different GFP versions were tested for protein production in CFPS. The left panel shows the architecture or plasmids with the mutation that each GFP variant carries. The final fluorescence of CFPS reactions is shown on the right. Data represents mean values ± standard deviation of three independent experiments.

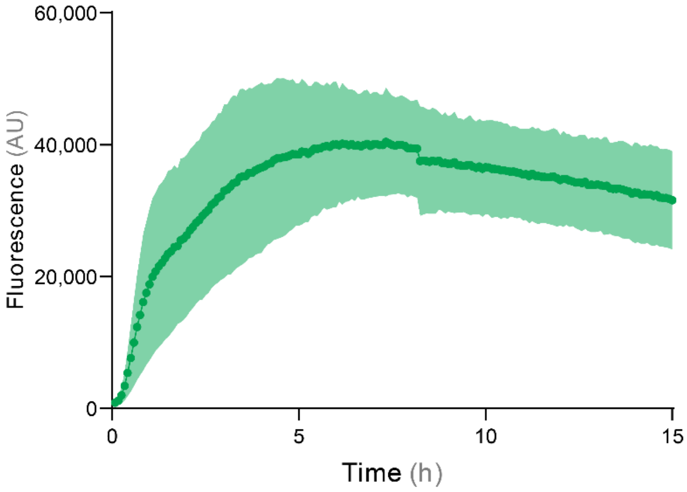


**Figure S3 |** The time-course assessment of the CFPS reaction with the lysate from strain BL21 Δ*endA*-ΔAA. Data represents mean values ± standard deviation of three independent experiments.

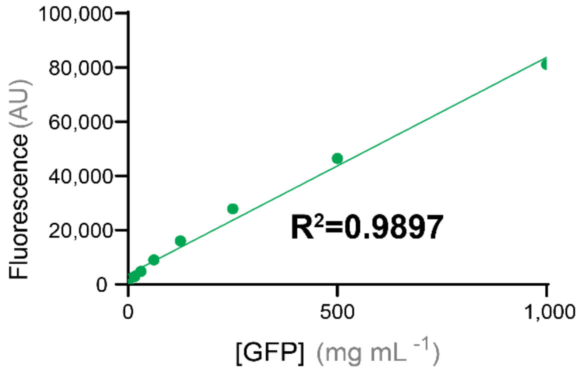


**Figure S4 |** GFP calibration curve generated using Abcam GFP Quantification kit. The curve was generated by diluting GFP in 1x PBS (pH 7.5) and fluorescence signals were measured using Biotek Synergy H 1 with excitation and emission at 485 and 528 nm, respectively.
